# Lamin B1 physically regulates neuronal migration by modulating nuclear deformability in the developing cortex

**DOI:** 10.1101/2025.10.22.683830

**Authors:** Minkyung Shin, Shunichi Ishida, Jinseon Yu, Misato Iwashita, Gyeong-un Jang, Pietro Cortelli, Elisa Giorgio, Ilaria Cani, Giulia Ramazzotti, Stefano Ratti, Daisuke Yoshino, Jong-Cheol Rah, Yohsuke Imai, Yoichi Kosodo

**Affiliations:** Neural Regeneration Laboratory, Korea Brain Research Institute (KBRI), Daegu, Republic of Korea; Graduate School of Engineering, Kobe University, Kobe, Japan; Neurophysiology Laboratory, KBRI, Daegu, Republic of Korea; Brain Sciences, Daegu Gyeongbuk Institute of Science & Technology (DGIST), Daegu, Republic of Korea; Alma Mater Studiorum, University of Bologna, Bologna, Italy; Department of Molecular Medicine, University of Pavia, Pavia, Italy; IRCCS Mondino Foundation, Neurogenetics Research Centre, Pavia, Italy; U.O.C. Clinica Neurologica Rete Metropolitana (NeuroMet), IRCCS Istituto delle Scienze Neurologiche di Bologna, Bologna, Italy; Cellular Signalling Laboratory, Department of Biomedical and Neuromotor Sciences (DIBINEM), University of Bologna, Bologna, Italy; Division of Advanced Applied Physics, Institute of Engineering, Tokyo University of Agriculture and Technology, Tokyo, Japan

**Author notes:** **Corresponding author:** Yoichi Kosodo, Korea Brain Research Institute (KBRI), 41062 Daegu, Republic of Korea.

**Keywords:** Lamin B1 (LB1), neuronal migration, cortical development, physical microenvironment, stiffness, autosomal dominant leukodystrophy (ADLD), human cerebral organoid (hCO)

## Abstract

Neuronal migration is a vital process that positions billions of neurons to create a functional brain. To navigate the constrained microenvironments within the cortex, precise control over the nuclear mechanics in migrating neurons is indispensable. Here, we show that Lamin B1 (LB1) regulates neuronal migration by modulating nuclear deformability. Excess LB1 in neurons halted migration without altering laminar identity or overall gene expressions *in vivo*, while *in vitro,* it elevated nuclear stiffness and impaired neuronal motility in confined spaces. Moreover, mispositioned neurons resulted in electrophysiological defects in the brain. Computational modeling predicted a temporal relationship between nuclear deformation and enhanced migration velocity, which was validated experimentally through live imaging. Notably, cerebral organoid assays using iPS cells established from patients with *LMNB1* duplication exhibited impaired neuronal migration in a human model. Collectively, these findings demonstrate that LB1 is a critical regulator of nuclear mechanics, ensuring the accurate spatiotemporal positioning of neurons.

## Introduction

The precise spatial and temporal organization of diverse neuronal populations is critical for optimal brain function. Particularly, most cortical excitatory neurons originate from neural progenitor cells (NPCs) located in the ventricular zone (VZ) of the developing brain, and migrate radially toward the pial surface to reach their designated layer positions (*1, 2*). Timely arrival at their final destinations is crucial for establishing functional neural circuits (*3*).

The distance and mode of migration vary depending on the neuronal subtype and the developmental timeline. For example, late-born excitatory neurons undergo longer radial migration to reach the superficial cortical layers (*4*). To accomplish such long-distance migration accurately, newborn neurons integrate extracellular guidance cues and undergo dynamic morphological transitions in the intermediate zone (IZ), switching from a multipolar to a bipolar shape (*5, 6*). This bipolar radial migration constitutes the primary mode of locomotion for excitatory neurons in the developing neocortex (*7*). The successful migration of bipolar neurons necessitates more than merely adhering to radial glial fibers; it also requires active cytoskeletal remodeling and nuclear translocation. Bipolar neurons extend and grasp a leading process along radial glial fibers, while their nuclei move through a combination of dynein-mediated pulling forces along microtubules and actomyosin-generated pushing forces from the rear (*8, 9*). This synergistic mechanism drives neuronal movement toward their destinations. In migrating neurons, the sequential extension of the leading process followed by delayed nuclear translocation results in a characteristic saltatory movement during locomotion.

While extensive research has focused on the cytoskeletal motors and signaling pathways involved in neuronal migration, the role of non-cell-autonomous factors is just beginning to receive attention (*10–12*). A major challenge lies in underscoring how neurons navigate through the confined spaces created by the extracellular matrix (ECM), axons, and particularly the cortical plate (CP), which is already occupied by earlier-born neurons. In such restricted spaces, the translocation of the nucleus, the largest and stiffest organelle, may emerge as a crucial physical factor for successful migration. Analogous circumstances have been noted in the migration of non-neuronal cells, including cancer and immune cells (*13–16*), yet the role of nuclear mechanics in neuronal migration warrants further exploration.

The nucleus is enclosed by a double-membrane structure consisting of an outer and inner nuclear membrane. Just beneath the inner nuclear membrane lies the nuclear lamina, a dense network of Lamins (type V intermediate filaments). This network connects to the linker of nucleoskeleton and cytoskeleton (LINC) complex, facilitating the mechanical coupling of the nucleus to cytoskeletal components, such as microtubules and actin filaments (*17*). Lamin-family proteins play a central role in regulating nuclear shape and mechanical properties (*18*). In mammalian cells, the nuclear lamina consists of A-type Lamins (Lamin A and C), produced through alternative splicing of the *LMNA* gene, and B-type Lamins (Lamin B1 (LB1) and B2 (LB2)), encoded by the *LMNB1* and *LMNB2* genes, respectively (*19*). Numerous studies have demonstrated that nuclear stiffness is modulated in a Lamin A and C-dependent manner (*20–22*).

In the developing brain, however, Lamin A and C are barely expressed, while Lamin B members predominate (*23*). Studies using knockout mice revealed that LB1 and LB2 are essential for proper brain development, although their functions are not fully redundant. Loss-of-function mutations in *Lmnb1* and *Lmnb2* both result in impaired neuronal migration, leading to a markedly reduced number of neurons and disorganized cortical layers. *Lmnb1*^Δ/Δ^ mice exhibit nuclear blebs accompanied by an asymmetric distribution of LB2, whereas *Lmnb2*□/□ mice display elongated nuclei with a uniform distribution of LB1 (*24–26*). These findings highlight the critical role of Lamin Bs in brain development. Nevertheless, the mechanisms governing the nuclear deformation and navigation of migrating neurons through the densely packed and confined extracellular environments of the developing cortex remain poorly understood. Thus, elucidating the contribution of nuclear mechanics to this process is essential for a comprehensive understanding of neuronal migration during corticogenesis.

Here, we present evidence that LB1 regulates neuronal migration by modulating the physical properties of the nucleus. In newborn neurons, LB1 overexpression halted migration without altering laminar identity or overall gene expression, leading to rounder nuclei and increased stiffness. Computational modeling suggested that elevated nuclear stiffness alone can impair neuronal migration and that nuclear deformation is temporally linked to enhanced migration, predictions validated through *in*□*vivo* live imaging. Of particular significance is its link to autosomal dominant leukodystrophy (ADLD) (*27, 28*), a rare brain disease caused by LB1 overexpression (*29–32*). We established patient-derived induced pluripotent stem cells (iPSCs) and identified a deficit in neuronal migration within human cerebral organoids (hCOs), underscoring the potential clinical relevance of our findings. Overall, this study documents that excessive LB1 renders the nucleus overly rigid, surpassing its deformable threshold and preventing neurons from navigating through densely packed tissue, a critical mechanism for accurate neuronal positioning and function in the brain.

## Results

### LB1 overexpression impairs neuronal migration without altering laminar identity or global transcriptional profile

While Lamin A/C is minimally expressed during mammalian neocortical development, LB1 is ubiquitously expressed across various neural cell types in both developing mouse and human brains (Fig. S1), as previously reported (*23*). To better understand the potential role of LB1 in neuronal migration, we overexpressed either mouse *Lmnb1* (mLB1) or human *LMNB1* (hLB1) together with enhanced green fluorescent protein (EGFP) in NPCs by *in utero* electroporation (IUE) at embryonic day (E) 15.5, a mid-to-late neurogenic stage when post-mitotic neurons travel long distances (Fig. 1A). Four days after electroporation (Postnatal day 0, P0), the majority of EGFP□ neurons in control pups actively migrated from the VZ toward the CP across the neocortex (Fig. 1B and C; P0). At P0, 50□% of EGFP□ migrating neurons were located in the IZ, defined as the area beneath the NeuN^+^ CP, in control pups, compared to 70□% in hLB1-overexpressing (OE) and 80□% mLB1-OE pups (Fig.□1D; P0). This result suggests that elevated LB1 delays radial neuronal migration. At P7, both hLB1- and mLB1-OE neurons were ectopically positioned within the CP and beneath it in the IZ (Fig.□1B–D; P7). This mislocalization persisted through P14 and was still evident 6 months after electroporation, indicating a long-lasting migration defect (Fig.□1B–D; P14 and 6M). Because of the inside-out pattern of cortical development, neurons with delayed migration arrived later than controls and ultimately settled in more superficial positions (*33*).

**Fig. 1.**
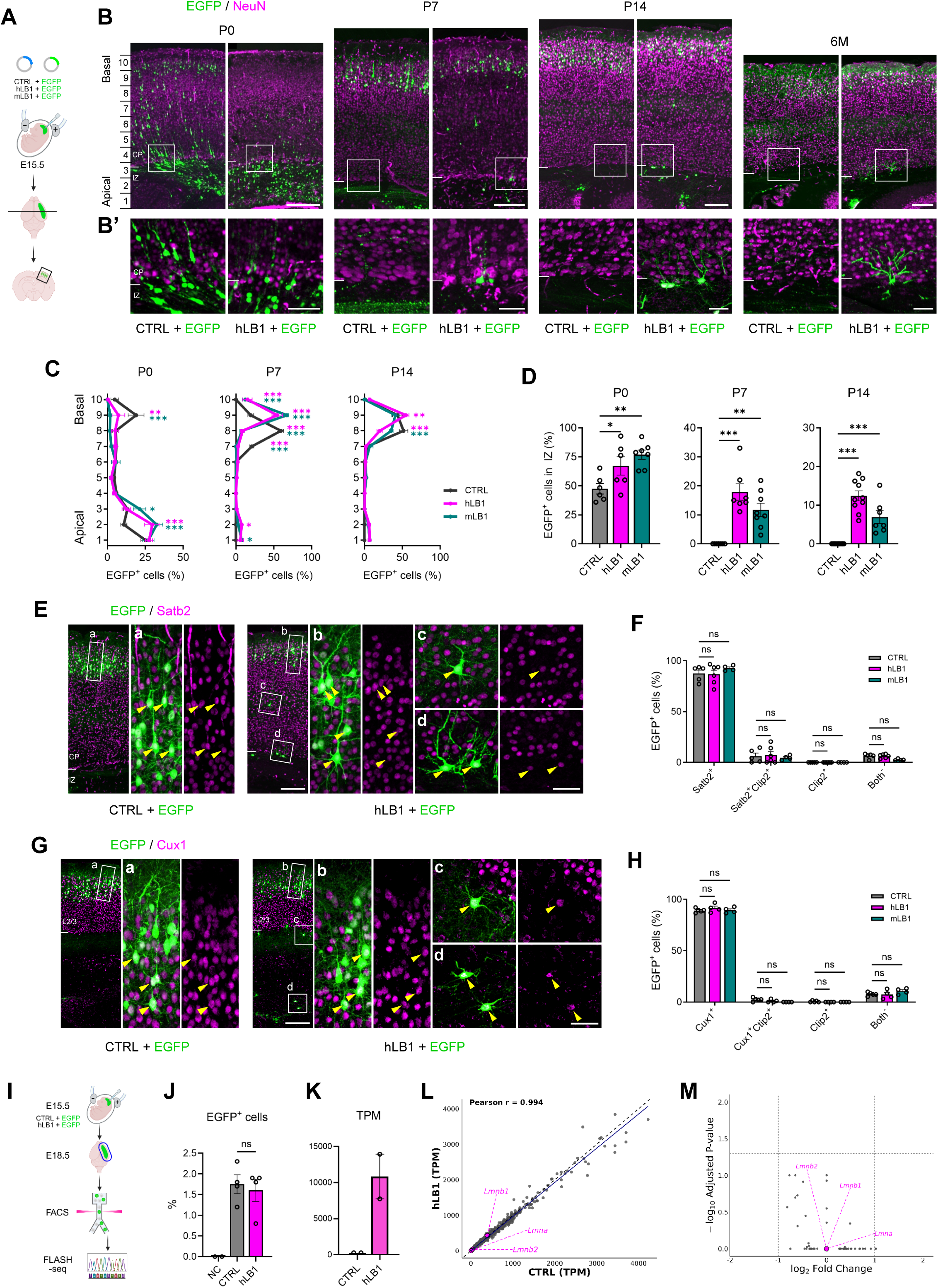
LB1-OE inhibits neuronal migration in the developing cortex. **(A)** Schematic of the IUE method. Created with BioRender.com. **(B)** The neocortex of E15.5 mice was electroporated with control (CTRL), human *LMNB1* (hLB1), or mouse *Lmnb1* (mLB1) plasmids with pCAG-EGFP, then fixed at P0, P7, P14, or at 6 months (6M). Coronal brain sections were stained with NeuN antibody (magenta). Bin numbers 1 to 10 are indicated along the apical-to-basal axis. (B’) Enlarged views of the boxed regions in (B) show EGFP+ (green) and NeuN+ neurons (magenta) in the IZ and CP. **(C)** Quantification of the laminar distribution of EGFP+ cells across bins 1 to 10, dividing the cortical wall from the VZ to the pial surface into ten equal bins (n = 6–7 brains per group). Counted cells for CTRL, hLB1, and mLB1 were 496, 295, and 387 (E15.5–P0); 219, 240, and 310 (E15.5–P7); 456, 451, and 441 (E15.5–P14), respectively. Statistical analysis was performed using two-way ANOVA followed by Dunnett’s multiple comparisons test. Error bars represent the mean ± SEM. \**P* < 0.05, \*\**P* < 0.01, \*\*\**P* < 0.001. **(D)** Quantification of EGFP+ cells in the IZ (n = 6–7 brains per group). Counted cells for CTRL, hLB1, and mLB1 were 496, 295, and 387 (E15.5–P0); 271, 233, and 290 (E15.5–P7); 362, 367, and 441 (E15.5–P14), respectively. Statistical analysis was performed using one-way ANOVA followed by Dunnett’s multiple comparisons test. Error bars represent the mean ± SEM. \**P* < 0.05, \*\**P* < 0.01, \*\*\**P* < 0.001. **(E)** Coronal sections at P14 were stained with Satb2 antibody (magenta). The white boxes indicate regions shown at higher magnification. Yellow arrowheads indicate EGFP+Satb2+ cells. **(F)** Quantification of EGFP+ cells expressing Satb2 and/or Ctip2 (n = 4–6 brains per group). Counted cells for CTRL, hLB1, and mLB1 were 287, 298, and 258 (E15.5–P14), respectively. Statistical analysis was performed using two-way ANOVA followed by Dunnett’s multiple comparisons test. Error bars represent the mean ± SEM. **(G)** Coronal sections at P14 were stained with Cux1 antibody (magenta). The white boxes indicate regions shown at higher magnification. Yellow arrowheads indicate EGFP+Cux1+ cells. **(H)** Quantification of EGFP+ cells expressing Cux1 and/or Ctip2 (n = 4–5 brains per group). Counted cells for CTRL, hLB1, and mLB1 were 332, 269, and 369 (E15.5–P14), respectively. Statistical analysis was performed using two-way ANOVA followed by Dunnett’s multiple comparisons test. Error bars represent the mean ± SEM. **(I)** Schematic of the FLASH-seq workflow. EGFP+ cells were collected 3 day after IUE by FACS and profiled using FLASH-seq. **(J)** Fraction of FACS-sorted EGFP+ cells from non-electroporated (NC), CTRL, and hLB1 brains collected at E18.5 (n = 4 mice per group). Statistical analysis was performed using Student’s *t*-test. **(K)** Quantification of human *LMNB1* transcripts from FLASH-seq data in FACS-sorted EGFP+ cells electroporated with either CTRL or hLB1. **(L)** Transcriptomic correlation of mean TPM values between CTRL and hLB1 groups. Each dot represents the mean TPM of a single gene (n = 24418). The fitted linear regression line (solid blue) and nuclear lamina genes (*Lmnb1*, *Lmnb2*, *Lmna*) (magenta) are highlighted. Genes with a TPM > 5000 (*Tuba1a*, *Actb*, *Fabp7*) were excluded. **(M)** Volcano plot of differential expression for CTRL versus hLB1, with log₂ fold change on the x axis and −log₁₀ adjusted *P* value on the y axis. Dashed lines indicate thresholds (|log₂FC| > 1; FDR < 0.05), with Lmnb1, Lmnb2, and Lmna are annotated (magenta). Scale bars: 200 µm (B, E, and G); 50 µm (B’, Ed, and Gd).

To examine whether LB1-OE affects neuronal subtypes, we conducted immunostaining using Ctip2 to label layer 5 (L5) cortical neurons, along with either Satb2 or Cux1 to indicate layer 2/3 (L2/3) cortical neurons. The majority (> 90%) of EGFP^+^ neurons across all conditions were positive for Satb2 or Cux1, while negative for Ctip2 (Fig. 1E–H). Moreover, the ectopic neurons induced by LB1-OE remained as Satb2^+^ and Cux1^+^ neurons (Fig. 1E and G). These results indicate that LB1-OE does not alter the laminar identity of cortical neurons.

The migration impairment induced by hLB1 overexpression may result from transcriptional dysregulation of specific gene sets. To investigate this, we isolated EGFP□ cells by fluorescence-activated cell sorting (FACS) at E18.5, a stage when active migration is ongoing, followed by RNA sequencing (Fig. 1I). The proportion of confidently isolated EGFP□ cells by FACS was approximately 1.5% across both conditions, with no significant differences (Fig. 1J and S2). Robust overexpression of *LMNB1* was confirmed by increased transcript abundance (Fig. 1K). Notably, endogenous *Lmnb1* transcripts were detected at higher levels than *Lmna* or *Lmnb2* in migrating neurons (Fig. 1L). Nevertheless, differential expression analysis revealed no significantly regulated mouse genes with an absolute log□ fold change of ±1 and a false discovery rate (FDR) < 0.05 (n = 24421 genes detected; Fig. 1M), suggesting that hLB1-OE in neurons does not lead to broad changes in gene expression. In line with the previous study, which found that a 50% knockdown of *LMNA* in cultured cells resulted in minimal proteomic changes (*34*), fluctuations in Lamin levels may have little effect on short-term expression changes. Together, LB1-OE interferes with neuronal migration across various developmental stages, without altering laminar identity or inducing global transcriptional changes.

### LB1 protein levels regulate nuclear deformability during neuronal migration

We next investigated whether variations in LB1 protein levels, which serve as a structural component of the nuclear lamina, could affect the nuclear morphology of neurons that failed to migrate to the CP. We performed immunostaining for LB1 in P0 pups to assess EGFP□ migrating neurons located in the IZ and the CP (Fig. 2A and B). Nuclear morphology was quantified by outlining the LB1-stained nuclei and measuring several parameters, including perimeter, area, circularity, and aspect ratio (AR) (Fig. 2C). A circularity value close to 1 indicates a more rounded shape, while the AR, defined as the ratio of the vertical to the horizontal axis, reflects the elongation of the nucleus along the migratory pathway. Compared to the control, LB1-OE neurons exhibited more rounded nuclei in both the IZ and the CP, showing increased circularity and decreased AR in the IZ, with even greater changes in the CP (Fig. 2D–G). Correlation analysis further revealed a significant relationship between LB1 protein intensity and nuclear circularity, indicating that elevated LB1 levels directly impair the nuclear deformability required for migration (Fig 2H). We also examined the leading processes of EGFP□ neurons located in the CP and found that both the process length and the process-to-total cell length (P/T) ratio increased in LB1-OE neurons. Additionally, there was a reduction in process thickness (Fig. 2I–L), which could reflect the morphological alteration caused by impeded somal translocation (*35, 36*).

**Fig. 2.**
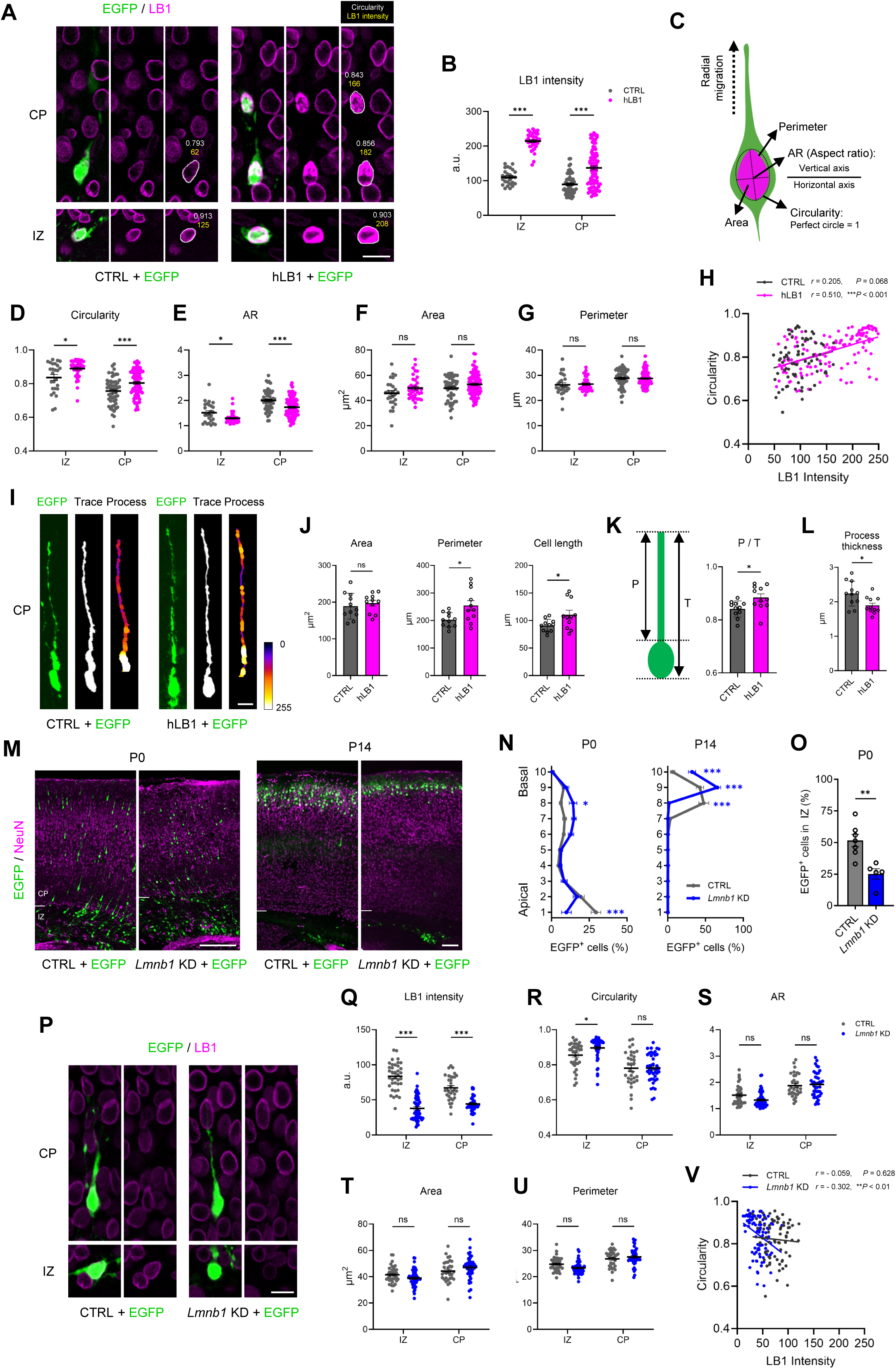
Nuclear deformability correlates with LB1 protein levels. **(A)** The neocortex of E15.5 mice was electroporated with CTRL or hLB1 plasmids along with pCAG-EGFP, then fixed at P0. Coronal brain sections were stained with LB1 antibody (magenta). The circularity values (white text) and LB1 protein intensity (yellow text) are indicated for each nucleus of EGFP+ cells. **(B)** Quantification of LB1 protein intensity in EGFP+ cells (n = 3–4 brains per group). Counted cells for CTRL and hLB1 were 26 and 36 (IZ) and 54 and 89 (CP), respectively. Statistical analysis was performed using two-way ANOVA followed by Šidák’s multiple comparisons test. Error bars represent the mean ± SEM. \*\*\**P* < 0.001. **(C)** Schematic of nuclear morphology analysis, including perimeter, AR, circularity, and area. The dotted black arrow indicates the direction of radial migration. (**D** to **G**) Quantification of nuclear morphology parameters in the same EGFP⁺ cells analyzed in (B) (n = 3–4 brains per group). Statistical analysis was performed using two-way ANOVA followed by Šidák’s multiple comparisons test. Error bars represent the mean ± SEM. \**P* < 0.05, \*\**P* < 0.01, \*\*\**P* < 0.001. **(H)** Pearson correlation analysis between LB1 protein intensity and nuclear circularity in each group. The fitted linear regression lines for each group are presented (black for CTRL and magenta for hLB1). A significant positive correlation was identified in hLB1 (*r* = 0.510, *P* < 0.001; n = 125 cells), but not in CTRL (*r* = 0.205, *P* = 0.068; n = 80 cells). **(I)** Representative tracing of an EGFP+ cells in the CP from CTRL and hLB1 groups, with pseudocoloring to indicate process thickness along the leading process. Color scale: 8-bit pixel values (0–255), where higher values represent thicker processes. **(J)** Quantification of cell morphology analysis including area (left), perimeter (middle), and total cell length (right) of migrating neurons in the CP based on EGFP fluorescence (n = 11 cells per group). Statistical analysis was performed using Student’s *t*-test. Error bars represent the mean ± SEM. \**P* < 0.05. **(K)** Schematic (left) and quantification (right) of the leading process-to-total cell length ratio (P/T), defined as the leading process length divided by the total cell length (n = 11 cells per group). Statistical analysis was performed using Student’s *t*-test. Error bars represent the mean ± SEM. \**P* < 0.05. **(L)** Mean process thickness measured along the entire length of the leading process (n = 11 cells per group). Statistical analysis was performed using Student’s *t*-test. Error bars represent the mean ± SEM. \**P* < 0.05. **(M)** The neocortex of E15.5 mice was electroporated with either scrambled control (CTRL) or Lmnb1-targeting shRNA (*Lmnb1* KD) plasmids with pCAG-EGFP, then fixed at P0 or P14. Coronal brain sections were stained with NeuN antibody (magenta). **(N)** Quantification of the laminar distribution of EGFP+ cells across bins 1 to 10, where the cortical wall from the VZ to the pial surface was divided into ten equal bins (n = 3–5 brains per group). Counted cells for CTRL and Lmnb1 KD were 973 and 509 (E15.5–P0); 565 and 332 (E15.5–P14), respectively. Statistical analysis was performed using two-way ANOVA followed by Šidák’s multiple comparisons test. Error bars represent the mean ± SEM. \**P* < 0.05, \*\*\**P* < 0.001. **(O)** Quantification of EGFP+ cells in the IZ at P0 (n = 5–7 brains per group). Counted cells for CTRL and *Lmnb1* KD were 504 and 249, respectively. Statistical analysis was performed using Student’s *t*-test. Error bars represent the mean ± SEM. \*\**P* < 0.01. **(P)** The neocortex of E15.5 mice was electroporated with CTRL or *Lmnb1* KD plasmids with pCAG-EGFP, then fixed at E18.5. Coronal brain sections were stained with LB1 antibody (magenta). (**Q** to **U**) Quantification of LB1 protein intensity and nuclear morphology parameters in EGFP+ cells (n = 3 brains per group). Counted cells for CTRL and *Lmnb1* KD were 37 and 51 (IZ) and 32 and 43 (CP), respectively. Statistical analysis was performed using two-way ANOVA followed by Šidák’s multiple comparisons test. Error bars represent the mean ± SEM. \**P* < 0.05, \*\*\**P* < 0.001. (**V**) Pearson correlation analysis between LB1 protein intensity and nuclear circularity in each group. The fitted linear regression lines for each group are presented (black for CTRL and blue for Lmnb1 KD). A significant positive correlation was identified in *Lmnb1* KD (*r* = - 0.302, *P* < 0.01; n = 94 cells), but not in CTRL (*r* = - 0.059, *P* = 0.628; n = 69 cells). Scale bars: 10 µm (A, I, and P); 150 µm (M).

Subsequently, we conducted a loss-of-function study using shRNA to target mouse *Lmnb1* (Fig. 2M–V and S3). In the control group, approximately 16% of EGFP□ neurons reached the upper bins of the CP, while this proportion increased to 25% in the *Lmnb1* knockdown (KD) group at P0 (Fig. 2M and N, P0; bins 8–10 combined). Consistently, 50□% of EGFP□ neurons remained in the IZ in control pups, whereas this proportion diminished to 25□% in the *Lmnb1* KD group (Fig.□2O). These results suggest that a reduction in LB1 expression enhances neuronal migration efficiency. At P14, *Lmnb1* KD brains exhibited an increased accumulation of EGFP□ neurons in the outermost layers of the CP compared to controls (Fig. 2M, N, and S4), possibly because phospho-ERK failed to translocate into the nucleus, a process required for dendrite development (*37*). Three days after electroporation (E18.5), *Lmnb1* KD reduced LB1 protein levels in EGFP^+^ migrating neurons in both the IZ and the CP, but did not affect nuclear circularity, AR, area, and perimeter in the CP. Conversely, in the IZ, the nuclei became rounder, showing higher circularity than controls, which indicates a mild loss of elongation (Fig. 2P–U). Previous studies reported that *Lmnb1* deficiency can lead to nuclear rupture and structural instability (*24, 38*). Based on these reports, we reasoned that neurons with markedly reduced LB1 may similarly influence nuclear shape during neuronal migration (Fig. 2V).

Collectively, these results demonstrate that LB1 levels bidirectionally regulate nuclear deformability and neuronal migration. Increased expression of LB1 impairs deformability and slows migration, while reduced expression enhances nuclear plasticity and facilitates migration.

### LB1 overexpression increases nuclear stiffness and restricts migration through confined environments

The correlation observed between nuclear deformability and the positioning of neurons in the developing brain prompted us to explore whether variations in the physical properties of the nucleus could influence neuronal migration. Lamin A/C plays a crucial role in regulating nuclear stiffness in specific cell types (*19*). Here, we investigated whether the level of LB1 impacts nuclear stiffness in neurons. We cultured primary cells dissociated from E18.5 cortex after introducing plasmids via IUE at E15.5. Twenty-four hours after plating, we quantified the stiffness in the cell bodies of EGFP□ cells using AFM with a 20□µm bead-tipped cantilever (*39*). The hLB1-OE cells showed elevated LB1 protein levels and increased stiffness compared to controls (Fig.□3A–D). Given that the nucleus occupies a large volume within the cell body and that LB1 protein localizes exclusively on the nuclear lamina, the differences in stiffness between the hLB1-OE and control cells are likely attributed to changes in nuclear stiffness.

Next, to test whether increased nuclear stiffness affects cell passage through confined spaces, we conducted an 8□µm-pore transwell assay (Fig. 3E) (*11*). Forty hours after seeding primary neurons on the upper surface (TOP) of the transwell membranes, we observed that the fraction of EGFP^+^ hLB1-OE neurons reaching the bottom surface (BOT) was decreased compared to the control group (Fig. 3F and G). This result indicates that an excess level of LB1 reduces migratory capacity under physical confinement, suggesting that increased nuclear stiffness impeded somal translocation.

**Fig. 3.**
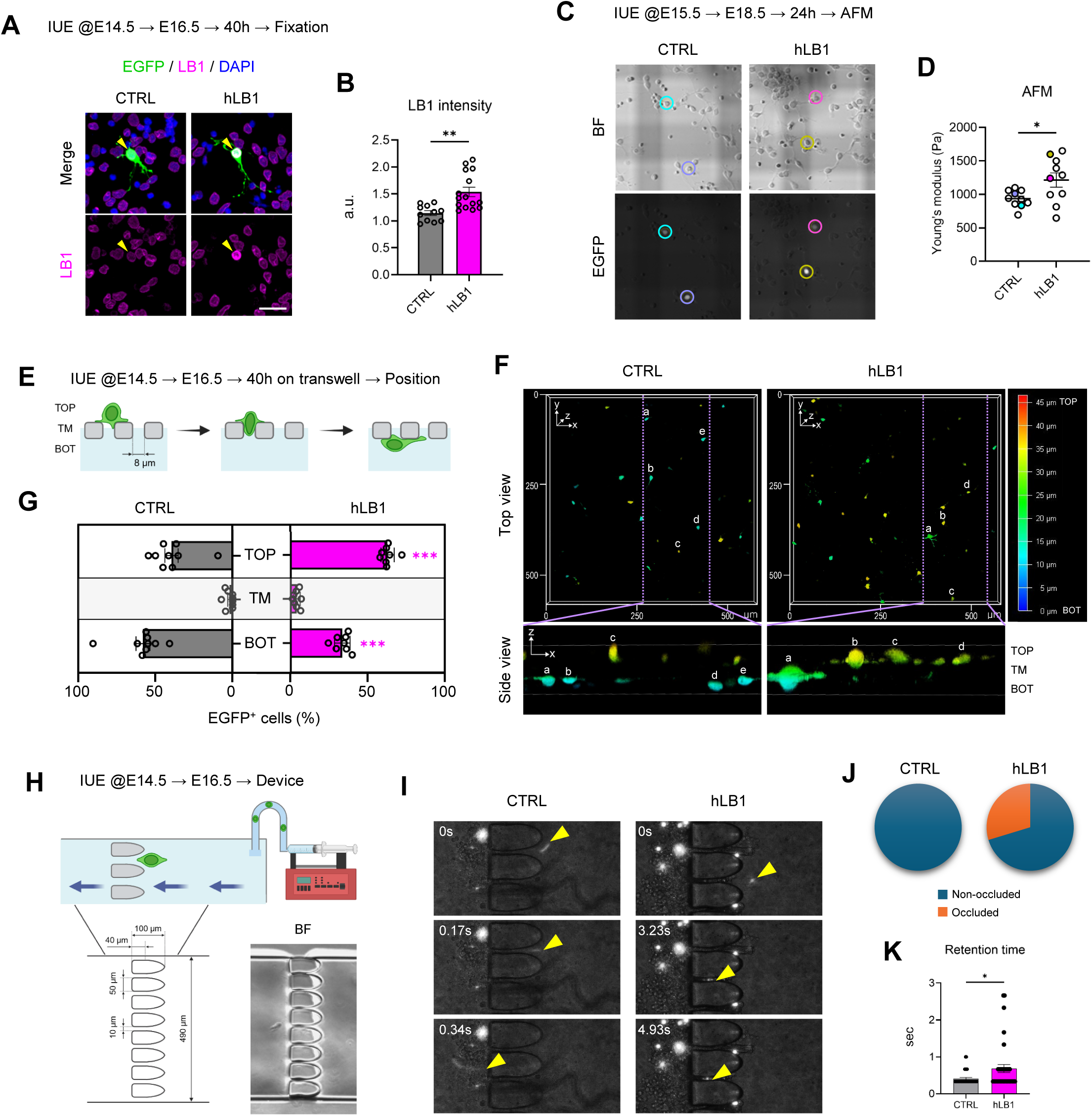
Increased stiffness due to LB1-OE impairs cell migration through narrow physical constraints. **(A)** The neocortex of E14.5 mice was electroporated with CTRL or hLB1 plasmids with pCAG-EGFP. At E16.5, the brains were dissected for primary culture. After 40 h, the cells were fixed and stained with LB1 antibody (magenta). Yellow arrowheads indicate EGFP+ cells. **(B)** Quantification of LB1 protein intensity in EGFP+ cells (n = 11–15 cells per group). Statistical analysis was performed using Student’s *t*-test. Error bars represent the mean ± SEM. \*\**P* < 0.01. **(C)** The neocortex of E15.5 mice was electroporated with CTRL or hLB1 plasmids with pCAG-EGFP. At E18.5, the brains were dissected for primary culture, and AFM measurements were performed on live cells after 24h. Colored circles indicate EGFP⁺ cells selected for AFM analysis, corresponding to the data presented in (D). **(D)** The stiffness of EGFP+ cells measured by AFM (n = 10 cells per group from n = 5–6 brains). Statistical analysis was performed using Student’s *t*-test. Error bars represent the mean ± SEM. \**P* < 0.05. **(E)** Schematic of the transwell migration assay. The neocortex of E14.5 mice was electroporated with CTRL or hLB1 plasmids with pCAG-EGFP. At E16.5, the brains were dissected, and the cells were placed in the upper chamber (TOP) of an 8-µm-pore transwell membrane (TM). After 40 h, the cells were fixed and their positions were quantified. **(F)** EGFP+ cells were imaged from both top and side views following the transwell assay. Letters (a–e) indicate the same EGFP⁺ cells in both views to indicate their relative positions across the transwell membrane. **(G)** Quantification of the proportion of EGFP+ cells on the top (TOP), transwell membrane (TM), and bottom (BOT) sides of the transwell membrane. Counted cells for CTRL and hLB1 were 282 and 626, respectively. Statistical analysis was performed using two-way ANOVA followed by Šidák’s multiple comparisons test. Error bars represent the mean ± SEM. \*\*\**P* < 0.001. **(H)** Schematic of the microfluidic device. The neocortex of E14.5 mice was electroporated with CTRL or hLB1 plasmids with pCAG-EGFP. At E16.5, the brains were dissected for the cell arrest assay. **(I)** Time-lapse imaging of EGFP+ cells migrating through narrow slits in the microfluidic chamber. Yellow arrowheads indicate EGFP+ cells entering, traversing, or being occluded within a slit. **(J)** Cells that remained in the slit for more than 5 seconds were classified as occluded, while those that passed through within 5 seconds were classified as non-occluded (n = 35–37 cells per group from n = 12–13 brains). **(K)** Retention time of EGFP+ cells, quantified from the non-occluded population (n = 26–35 cells per group from n = 12–13 brains). Statistical analysis was performed using Student’s *t*-test. Error bars represent the mean ± SEM. \**P* < 0.05. Scale bar: 20 µm (A).

To further elucidate the effects of nuclear stiffness on migration through confined spaces, we introduced EGFP□ cells into a microfluidic device with externally applied forces (Fig. 3H) (*15*). The device operated with a 1.0□µl/min flow, driving cells through eight constrictions separated by 10□µm gaps, to reconstitute *in*□*vivo* spatial restrictions (Fig. 3I). Approximately 30% of hLB1-OE cells failed to traverse the gaps within 5 seconds, leading to their classification as “occluded” (Fig. 3J). Among the cells that successfully crossed the gaps, the mean retention time for hLB1-OE cells was 1.7-fold longer than that of the control group (Fig. 3K).

Together, our findings from *in vitro* assays indicate that LB1-mediated stiffness reduces nuclear deformability, thereby restricting neuronal transit through confined spaces. This establishes a causal link between altered nuclear mechanics and impaired motility.

### Computational modeling predicts that a stiffer nucleus impedes cell motility

To explore how the mechanical properties of the nucleus affect neuronal migration, we developed a three-dimensional computational model that simulates a cell migrating through a densely packed environment. In this model, the cell was represented as a soft capsule with a stiffer core inside, mimicking a migrating neuron with a nucleus. The densely packed environment was modeled as immobile obstacles fixed in a viscous fluid to simulate cellular structures and their surrounding matrix in the CP. Both the cell and obstacles have a diameter of 10 µm, while the nucleus has a diameter of 6 µm (Fig.□4A).

In the simulation, we introduced two variable parameters: the spacing between the immobile obstacles (δ) and the stiffness of the nucleus within the cell (RGs). RGs is defined as the ratio of nuclear membrane stiffness to surface membrane stiffness, with the latter being a fixed value. We visualized the movement of the cell from both side and rear views to test values for δ and RGs (Fig. 4B and C). First, we assessed how variations in δ affected the cell’s ability to navigate through the immobile obstacles. Our *in vivo* measurements in the mouse CP indicated that the mean distance between the centers of three neighboring nuclei was approximately 7.27□µm (Fig. S5). We examined δ values of 6.8, 7, and 7.2□µm in the simulation and found that 7.2 µm was the optimal value (Fig. 4B, Movie S1). Second, we evaluated the effect of nuclear stiffness on cell migration by comparing RGs values of 2 (soft), 4, 8, and 16 (hard) (Fig. 4C, Movie S2). We selected RGs values of 4 and 8 for further simulation analysis.

**Fig. 4.**
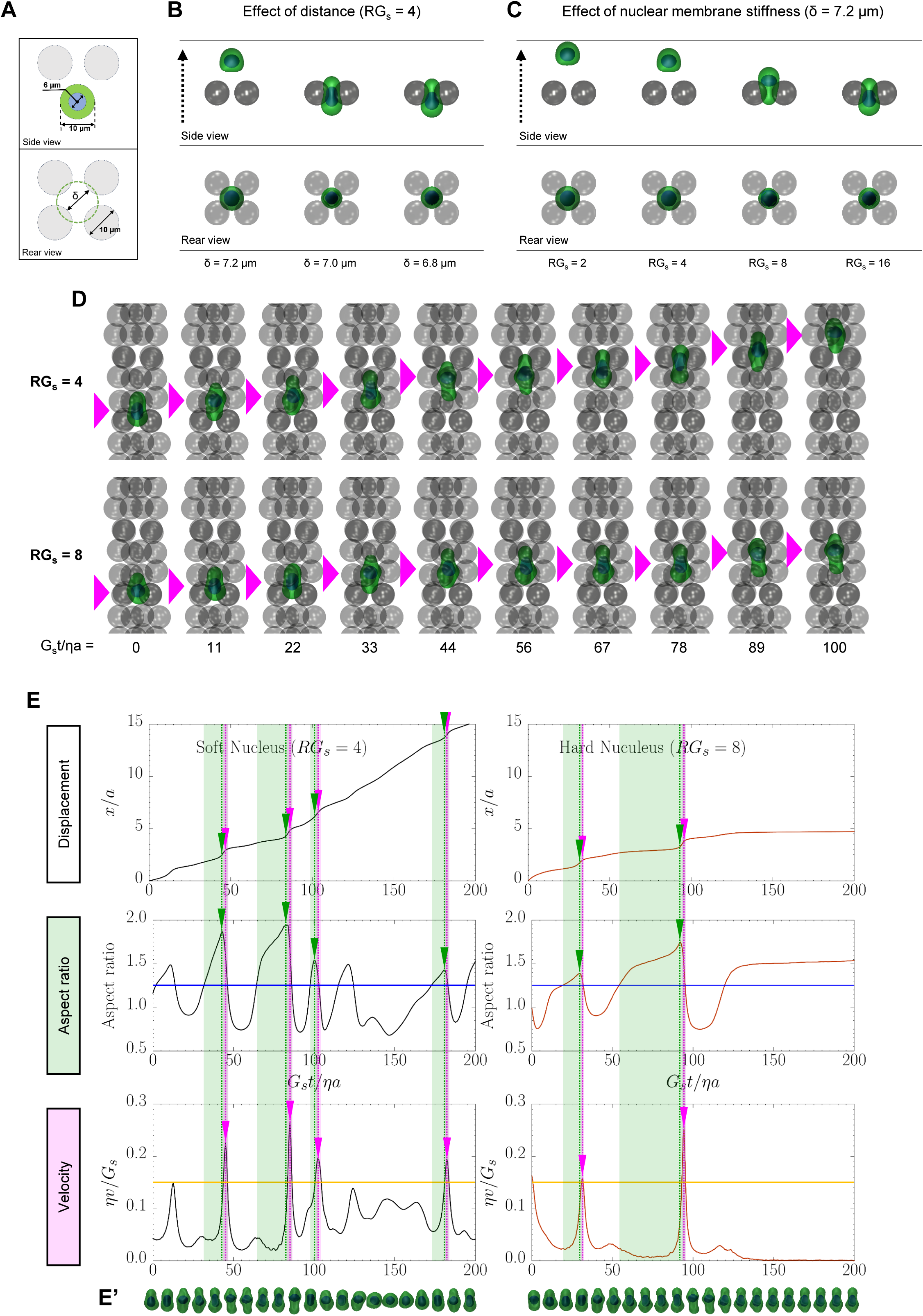
A simulation model of neuronal migration to analyze the effects of varied nuclear stiffness. **(A)** The design diagram of the simulation model, depicting both side (top) and rear (bottom) views. **(B)** The effect of the distance between immobile obstacles on neuronal migration at the same time point after the initiation of the simulation. The surrounding obstacles form a single layer. **(C)** The effect of nuclear membrane stiffness on neuronal migration. “RGs = 4” indicates that the nuclear membrane is 4 times stiffer than the cellular membrane. **(D)** Tracking of cell positions within an array of immobile obstacles under conditions of RGs = 4 (top) and 8 (bottom). The nuclear positions at each time point are indicated by magenta arrowheads. **(E)** Plots of displacement, AR, and velocity under conditions of RGs = 4 (left) and 8 (right) over time. Segments where the AR trace (blue) exceeds 1.25 are highlighted in green, with the corresponding peaks marked by green arrowheads. Segments where the velocity trace (orange) exceeds 0.15 are highlighted in magenta, with the corresponding peaks marked by magenta arrowheads. (E’) Tracking of cellular and nuclear shapes in each group.

Next, we performed simulations to assess how changes in nuclear stiffness impact vertical migration through a continuously confined environment that mimics the CP (Fig. 4D, Movie S3). Under softer conditions (RGs = 4), we observed that nuclear deformation, characterized by a higher AR, preceded each peak in cellular velocity (Fig. 4E, left). In the harder condition (RGs = 8), we recognized a similar trend at the early time point, where increased deformation correlated with higher velocities. However, after reaching a certain threshold, nuclear deformation stalled, resulting in a cessation of cellular migration (Fig. 4E, right). This stagnation appeared to stem from the inability of the stiff nucleus to deform and squeeze through narrow gaps, leading to mechanical entrapment between immobile obstacles. Our simulation results suggest a model in which the stiffness of the nuclear membrane modulates the AR, showing a considerable correlation with neuronal migratory behavior.

### LB1 overexpression decreases nuclear deformability and delays radial migration *in vivo*

Our computational model predicted that the stiffening of the nuclear membrane reduces nuclear deformability, as indicated by the fluctuations in the AR, ultimately migratory failure due to mechanical entrapment during obstacle traversal. To test whether these mechanical constraints affect neuronal migration in the developing brain, we performed live imaging *in vivo*. We simultaneously visualized nuclear and cellular morphology by co-electroporating nuclear-localizing H2B-TagRFP and cytoplasm-localizing EGFP. We initiated time-lapse imaging in the sliced brain 3 days after IUE at E15.5, a stage when radial migration is actively occurring. Images were captured every 5 minutes to track the dynamic behaviors of neuronal migration. To determine the initial positioning of EGFP□ neurons, we fixed littermate embryos and delineated cortical layers based on NeuN immunostaining (Fig. 5A). The majority of EGFP□ neurons were located in the IZ at the start of imaging, with ∼60% present in both the control and hLB1-OE groups (Fig. 5B).

**Fig. 5.**
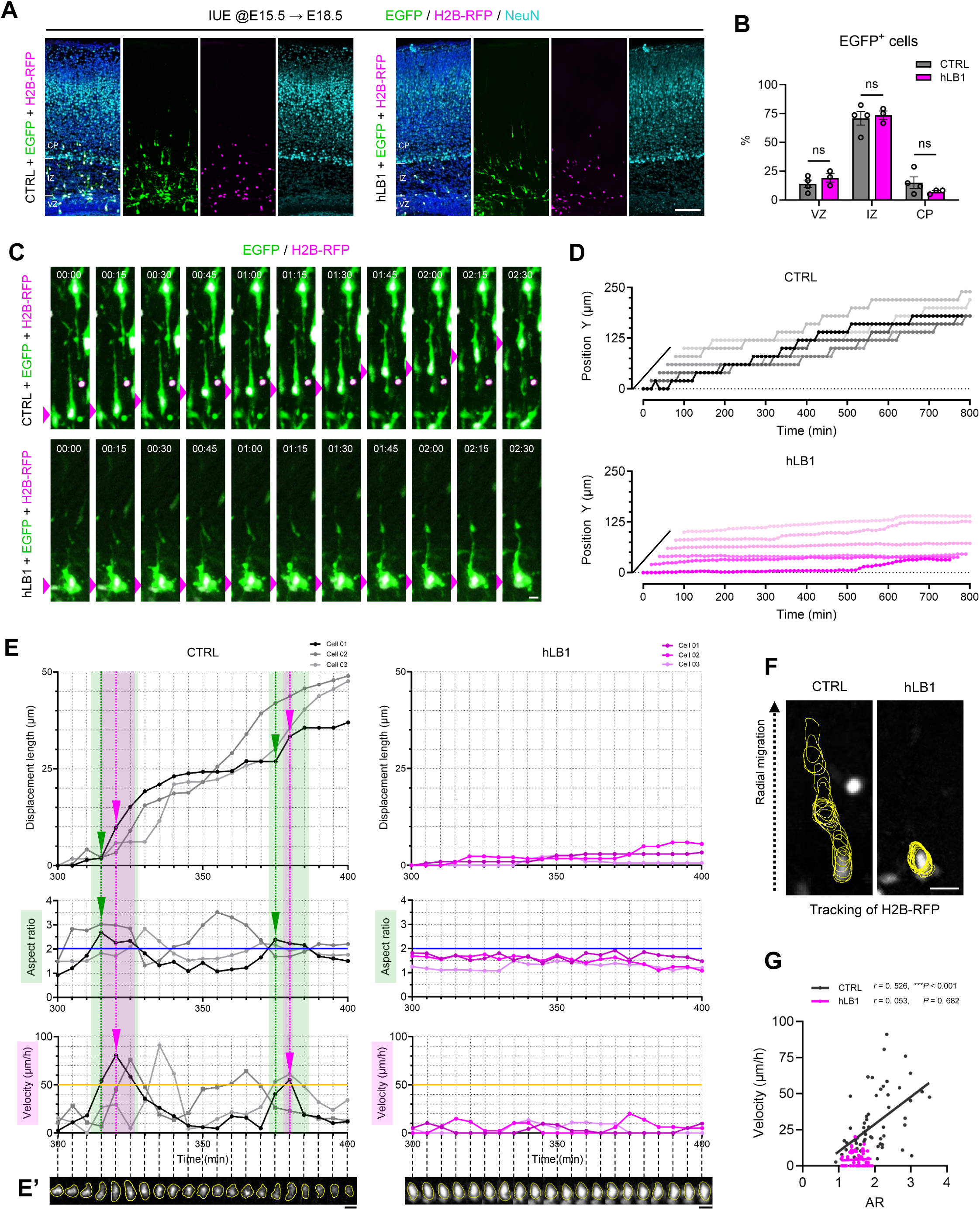
Time-lapse imaging to assess the correlation between neuronal migration and nuclear deformability in brain tissue. **(A)** The neocortex of E15.5 mice was electroporated with CTRL or hLB1 plasmids with pCAG-EGFP and pCAG-H2B-TagRFP, then fixed at E18.5, the starting stage of the time-lapse imaging in the brains of littermates. Coronal brain sections were stained with NeuN antibody (cyan). **(B)** Quantification of the relative positions of EGFP+ cells (n = 3–4 brains per group). Counted cells for CTRL and hLB1 were 148 and 265, respectively. Statistical analysis was performed using two-way ANOVA followed by Šidák’s multiple comparisons test. Error bars represent the mean ± SEM. **(C)** Cellular and nuclear morphology is visualized by EGFP and H2B-TagRFP signals in each group. Images were acquired at 5-minute intervals, with every third frame displayed here at 15-minute intervals. Elapsed time (h:min) is shown above each panel. The nuclear positions in each time point are indicated by magenta arrowheads. **(D)** Y-position plots of individual EGFP+ cells in the CTRL and hLB1 groups, showing the trajectories traced from the H2B-TagRFP nuclear signal. Each line represents the radial migration trajectory of a single cell (n = 6 cells per group). **(E)** Quantification of nuclear dynamics in individual EGFP+ cells at 5-minute intervals. The plots show displacement length (top), AR (middle), and velocity (bottom) for the CTRL (left) and hLB1 (right) conditions. Segments where the AR trace (blue) exceeds 2 are highlighted in green, with the corresponding peaks marked by green arrowheads. Segments where the velocity trace (orange) exceeds 50 µm/h are highlighted in magenta, with corresponding peaks marked by magenta arrowheads. (E’) Nuclear morphology is visualized by H2B-TagRFP signal in each group (Cell 01) at the corresponding time points. **(F)** The nuclear positions in EGFP+ cells tracked from the H2B-TagRFP signal (yellow lines), are imaged at 5-minute intervals over 20 frames. The dotted black arrow indicates the direction of radial migration. **(G)** Pearson correlation analysis between AR and velocity using data from the three cells shown in (E), measured at 5-minute intervals. The fitted linear regression lines for each group are presented (black for CTRL and magenta for hLB1). A significant positive correlation was identified in CTRL (*r* = 0.526, *P* < 0.001; n = 3 cells), but not in hLB1 (*r* = 0.053, *P* = 0.682; n = 3 cells). Scale bars: 100 µm (A); 10 µm (C, E’, and F).

Despite the initial similarity in spatial distribution, cell behaviors diverged over time; control neurons migrated upward toward the CP, while hLB1-OE neurons often remained immobile within the IZ (Fig. 5C). To further analyze these migration dynamics, we examined cell trajectories along the apical-basal (ventricle-pial) axis and found that the migratory patterns were markedly different between the CTRL and hLB1-OE groups. Control migrating neurons displayed saltatory movement, characterized by a “stop-and-go” pattern consistent with known neuronal behaviors (Fig. 5D) (*5, 6*). In contrast, hLB1-OE neurons failed to exhibit such coordinated progression.

To assess the relationship between nuclear shape and neuronal migration, we tracked H2B-TagRFP signals at 5-minute intervals over a 100-minute period. Our quantitative analysis revealed that□(i) nuclear deformation, indicated by an elevated AR, persisted longer than changes in nuclear velocity;□(ii) the maximum velocity was observed within 10 min following the peak of nuclear elongation, occasionally occurring simultaneously; and (iii) this behavior recurred within individual neurons at intervals of 30–60 min (Fig. 5E and S6). In contrast, hLB1-OE neurons exhibited limited changes in nuclear shape, with AR values consistently hovering around 1.5, resulting in minimal displacement throughout the imaging period (Fig. 5E and F). The average migration velocity for control neurons was approximately 27□µm/h, while hLB1-OE neurons only reached about 4□µm/h. These *in*□*vivo* observations were consistent with our computational simulations, supporting a time-dependent relationship between nuclear deformability and neuronal motility. Importantly, we found a positive correlation between migration velocity and nuclear elongation, reinforcing the notion that reduced nuclear deformability significantly limits cell motility (Fig. 5G). Overall, our findings demonstrate that LB1-induced nuclear stiffening disrupts the shape remodeling necessary for saltatory translocation, thereby delaying neuronal migration *in vivo*.

### Ectopic positioning of LB1-OE neurons alters intrinsic excitability

To examine the physiological consequences of the LB1-dependent migration defect, we conducted whole-cell patch-clamp recordings on EGFP□ neurons in the somatosensory cortex at P14–15. This developmental stage allows for layer-specific neuronal comparisons, and the intrinsic firing properties of cortical pyramidal neurons are approaching adult-like levels, although full electrophysiological maturation has not yet been achieved (*40, 41*). In control brains, EGFP□ neurons were predominantly localized to L2/3, consistent with the expected positions of normally migrated neurons, whereas hLB1-OE neurons were observed not only in L2/3 but also ectopically in cortical layers other than L2/3 and in the IZ (Fig. 6A and B).

**Fig. 6.**
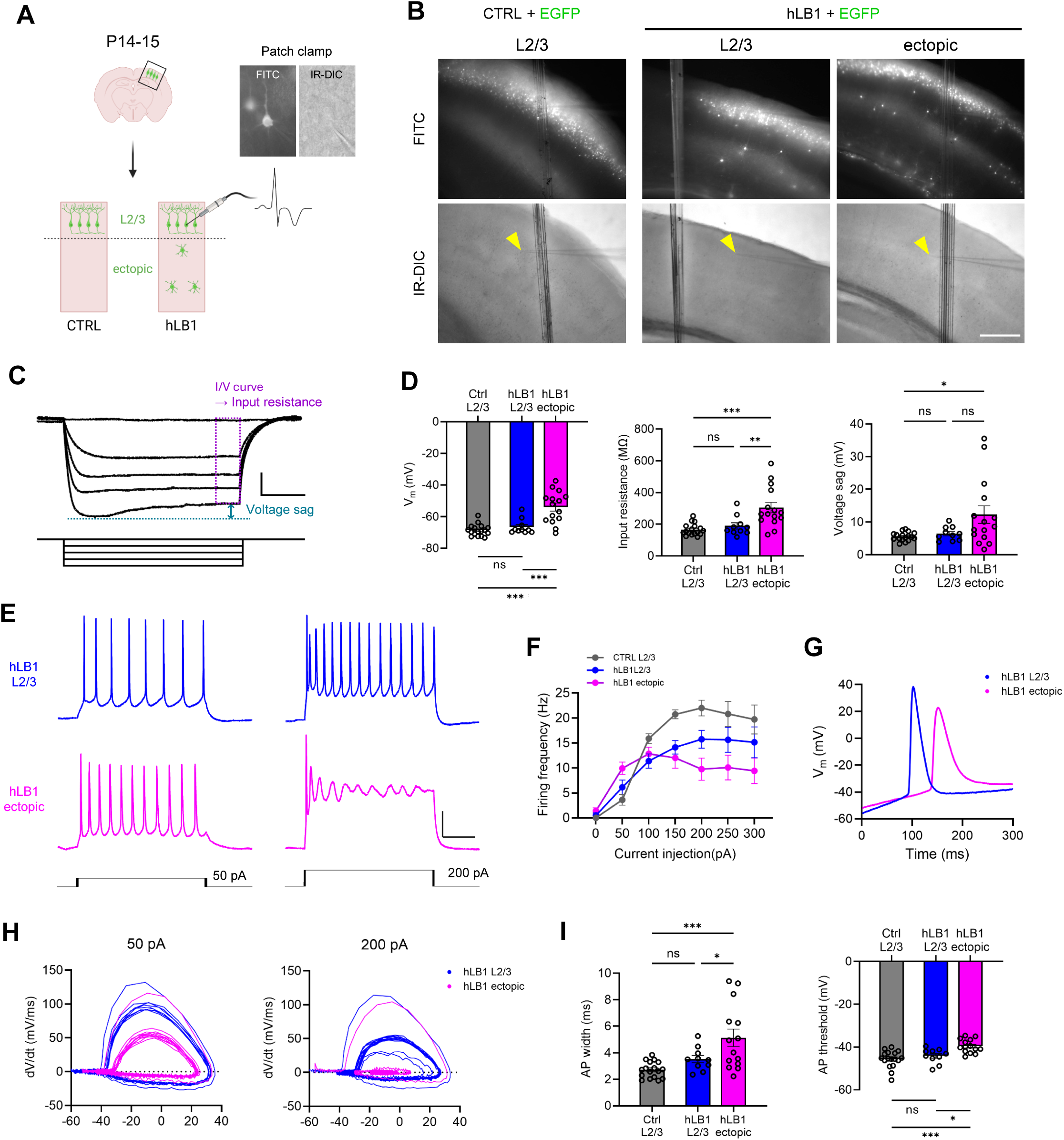
Ectopic positioning of hLB1-OE neurons alters intrinsic electrophysiological properties. **(A)** Schematic of whole-cell patch-clamp recordings. The neocortex of E15.5 mice was electroporated with CTRL or hLB1 plasmids with pCAG-EGFP. At P14-15, the brains were dissected for patch-clamp recordings. **(B)** Representative FITC and IR-DIC images show EGFP+ cells located in L2/3 or ectopic positions within primary somatosensory cortex. Yellow arrowheads indicate capillary tips used in patch-clamp recordings. **(C)** Representative voltage responses to hyperpolarizing and depolarizing current steps used to analyze passive membrane properties. **(D)** Quantification of resting membrane potential (Vm, left), input resistance (middle), and voltage sag (right) of EGFP+ neurons (n = 10–17 cells per group). **(E)** Representative traces showing AP firing in response to 600-ms current injections of 50 pA (left) and 200 pA (right) in hLB1 L2/3 (blue) and hLB1 ectopic (magenta) neurons (n = 10–17 cells per group). **(F)** Quantification of firing frequencies in response to 600-ms current injections (0–300 pA in 50 pA steps) in control L2/3 neurons (gray), hLB1 L2/3 neurons (blue), and hLB1 ectopic neurons (magenta). **(G)** Overlay of representative first AP traces evoked by 100 pA current injection, illustrating altered spike kinetics in hLB1 ectopic neurons. **(H)** Representative phase-plane plots (dV/dt vs. membrane potential) from hLB1 L2/3 (blue) and hLB1 ectopic (magenta) neurons in response to 50 pA (left) and 200 pA (right) current injections. **(I)** Quantification of AP properties across groups. AP width (left) was measured as the mean of multiple spikes evoked at each cell’s rheobase current. AP threshold (right) was determined from the first spike evoked by a 100 pA current injection (n = 10–17 cells per group). Statistical analysis was performed using one-way ANOVA followed by Tukey’s multiple comparison test. Error bars represent the mean ± SEM. \**P* < 0.05, \*\**P* < 0.01, \*\*\**P* < 0.001. Scale bars: 400 µm (B).

We assessed the intrinsic membrane properties of these neurons by applying a series of hyperpolarising steps (−200 to 0 pA, 50 pA increments; Fig. 6C). Intriguingly, the resting membrane potential, input resistance, and hyperpolarization-activated cation current-dependent voltage sag were indistinguishable between control and hLB1-OE neurons in L2/3 (Fig. 6D). In contrast, ectopic hLB1-OE neurons were significantly more depolarized, exhibited higher input resistance, and showed an enhanced voltage sag, hallmarks of electrophysiological immaturity in cortical neurons, as previously reported (*41, 42*) (Fig. 6D).

Depolarizing current steps (0–400□pA in 50□pA increments) revealed marked alterations in active membrane properties of hLB1-OE neurons. Unlike hLB1 L2/3 neurons, ectopic hLB1-OE neurons were unable to keep firing with increasing current injection, indicating reduced capacity to sustain repetitive spiking (Fig.□6E and F). In response to a 100□pA current injection, ectopic hLB1-OE neurons displayed a broader first action potential (AP) and a depolarized spike threshold (Fig.□6G). To further characterize these abnormalities in firing kinetics, we analyzed the rate of membrane potential change (dV/dt) during APs evoked by either rheobase current injection (∼50□pA) or a suprathreshold stimulus (200□pA), which reliably elicited spiking in most neurons. Phase-plane plots revealed reductions in both the maximum and minimum dV/dt in ectopic hLB1-OE neurons (Fig.□6H), and this impairment in repolarization became more pronounced at higher current intensities, indicating progressive impairment in AP repolarization. These alterations suggest slowed rising and repolarization phases, potentially due to immature or reduced activation of voltage-gated Na□ and K□ channels (*43*). Consistent with these kinetic changes, ectopic hLB1-OE neurons exhibited significantly increased spike width and more depolarized AP threshold compared to control and hLB1 L2/3 neurons (Fig.□6I).

These features, including prolonged spike width, depolarized threshold, and attenuated dV/dt peaks, are hallmarks of electrophysiologically immature cortical neurons. Collectively, these findings suggest that delayed neuronal maturation and altered AP kinetics contribute to the intrinsically immature excitability of ectopic hLB1-OE neurons. Despite expressing L2/3 markers such as Satb2 and Cux1 (Fig.□1E–H), these mispositioned neurons remain in an immature electrophysiological state, highlighting a mechanistic link between nuclear architecture, laminar misplacement, and AP kinetics.

### Cortical neurons induced from ADLD patient-derived iPSCs exhibit increased LB1 and stiffness

We further aimed to explore whether LB1-dependent nuclear stiffening affects neuronal migration in a human model. To achieve this, we employed a strategy to induce cortical neurons from iPSCs established from ADLD patient-derived cells. Using the Yamanaka factors (*44*), we reprogrammed dermal fibroblasts from 5 ADLD patients carrying an *LMNB1* duplication and 2 control individuals (Fig. 7A and S7A). The control individuals are genetically related to the ADLD patients but do not have the *LMNB1* duplication, making them suitable for comparative analysis. We confirmed that patient-derived fibroblasts exhibited elevated LB1 protein levels compared to the controls (Fig. 7B). Both control- and ADLD-iPSC lines displayed comparable expression levels of pluripotency genes and normal karyotypes (Fig. S7B–F). Consequently, these ADLD-iPSCs, which possess the *LMNB1* genetic duplication, provide a viable human model of LB1 overexpression.

**Fig. 7.**
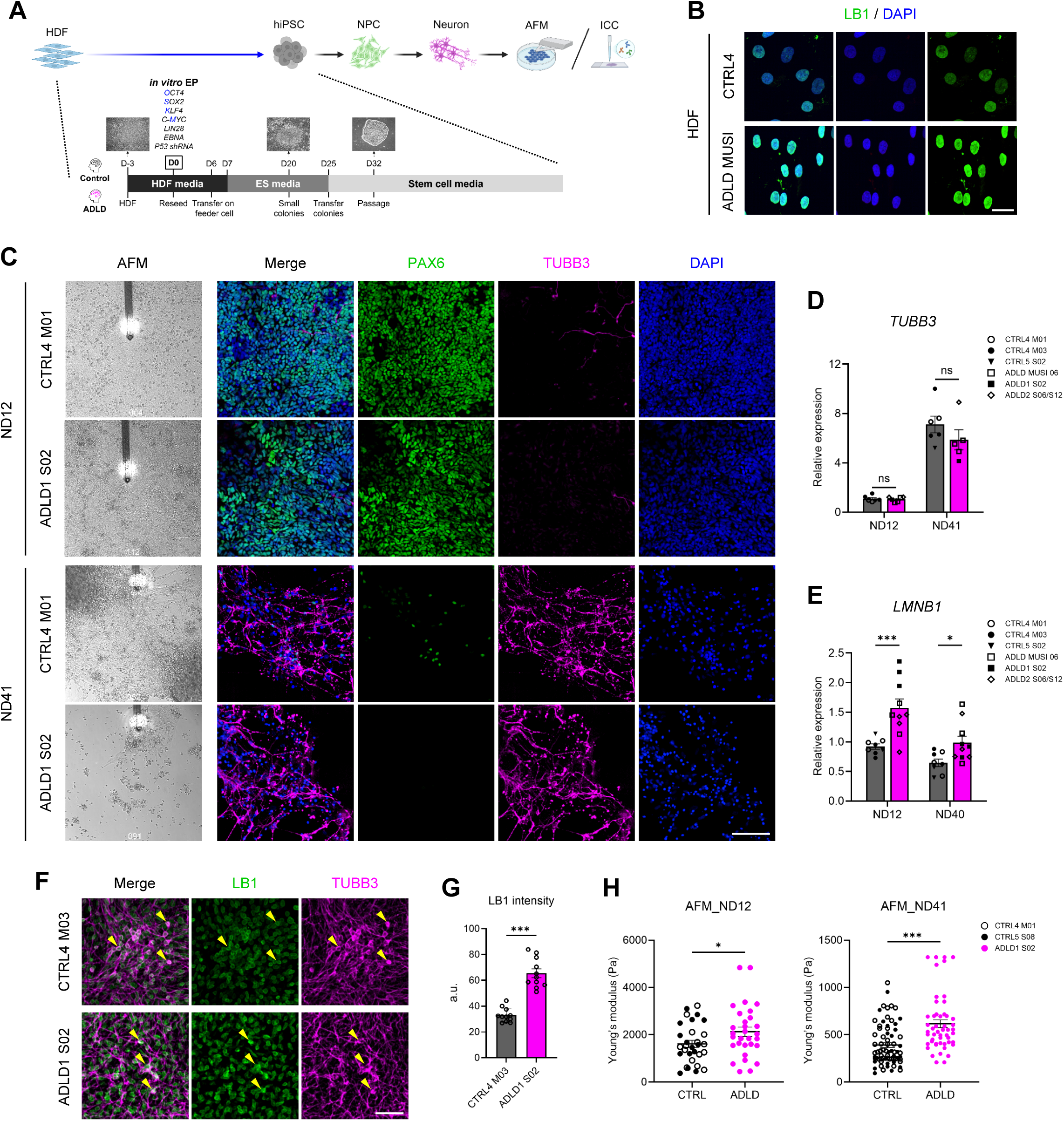
Establishment of ADLD patient-derived iPSCs and measurement of stiffness in NPCs and neurons. **(A)** Schematic of the generation of iPSCs and subsequent neural induction from CTRL and ADLD patient-derived cells. **(B)** Human dermal fibroblasts (HDFs) from CTRL and ADLD patients were stained with LB1 antibody (green). **(C)** Neural differentiation from CTRL and ADLD patient-derived iPSCs. At ND12 and ND41,the stiffness of live cells was measured using AFM. At the same time points, cells were fixed and stained with PAX6 (green) and TUBB3 (magenta) antibodies, and counterstained with DAPI (blue). (**D** and **E**) Quantification of relative mRNA expression levels of *TUBB3* and *LMNB1* at ND12 and ND40 measured by qPCR from two controls and three ADLD patient-derived cells. Each dot represents an independent experiment. Statistical analysis was performed using two-way ANOVA followed by Šidák’s multiple comparisons test. Error bars represent the mean ± SEM. \**P* < 0.05, \*\*\**P* < 0.001. **(F)** At ND40, cells were stained with LB1 (green) and TUBB3 (magenta) antibodies. Yellow arrowheads indicate TUBB3+ neurons. **(G)** Quantification of LB1 protein intensity in TUBB3+ neurons in (F) (n = 11 cells per group). Statistical analysis was performed using Student’s *t*-test. Error bars represent the mean ± SEM. \*\*\**P* < 0.001. **(H)** Quantification of the stiffness of CTRL and ADLD at ND12 and ND41. Counted cells for CTRL and ADLD were 30 and 30 (ND12) and 53 and 33 (ND41), respectively. Statistical analysis was performed using Student’s *t*-tests. Error bars represent the mean ± SEM. \**P* < 0.05, \*\*\**P* < 0.001. Scale bars: 30 µm (B);100 µm (C); 50 µm (F).

We conducted neural differentiation using iPSCs derived from both control and ADLD individuals (Fig. 7C). At neural differentiation day 12 (ND12), most of the cells expressed PAX6, indicating a predominant population of NPCs. At ND40–41, most cells exhibited a differentiated neuronal identity, as evidenced by their *TUBB3* expression. QPCR analysis of *TUBB3* expression revealed no significant differences between the control and patient-derived cells at either ND12 or ND41, suggesting that the overall efficiency of neural differentiation was comparable between the groups (Fig. 7D). Importantly, both *LMNB1* transcript levels were consistently elevated by 1.5 to 2-fold in ADLD cells compared to controls at both ND12 and ND40 (Fig. 7E). At ND40, LB1 protein levels in TUBB3^+^ neurons were about 2-fold higher in ADLD patient lines (Fig. 7F and G). Consistent with elevated LB1 both in transcriptional and translational levels, AFM measurements revealed increased stiffness in the ADLD patient-derived cells at both time points (Fig. 7H), similar to what was observed in the hLB1-OE mouse cells (Fig. 3). Collectively, we established a human *in*□*vitro* model that reflects the LB1-OE paradigm created in mice through IUE. In this system, both the NPCs and neurons showed elevated LB1 mRNA and protein levels, which were associated with increased nuclear stiffness.

### Newborn neurons derived from ADLD patients exhibit impaired radial migration in brain organoids

Using the iPSC lines, we investigated neuronal migration in hCOs, which provide a three-dimensional environment closely resembling the physical constraints of actual brain tissue. We examined the overall developmental status of the hCOs by measuring their size from day 10 (D10) to D70 in an ADLD-patient line (ADLD1 S02) and its isogenic controls (CTRL5 S08), revealing no significant differences at any time point (Fig. 8A–C). Consistent with results from two-dimensional cultures, the intensity of LB1 protein was 1.5-fold higher in ADLD-hCOs compared to the controls, whereas PAX6^+^ NPCs and CTIP2^+^ L5 neurons were similarly distributed in both groups (Fig. 8D-H).

**Fig. 8.**
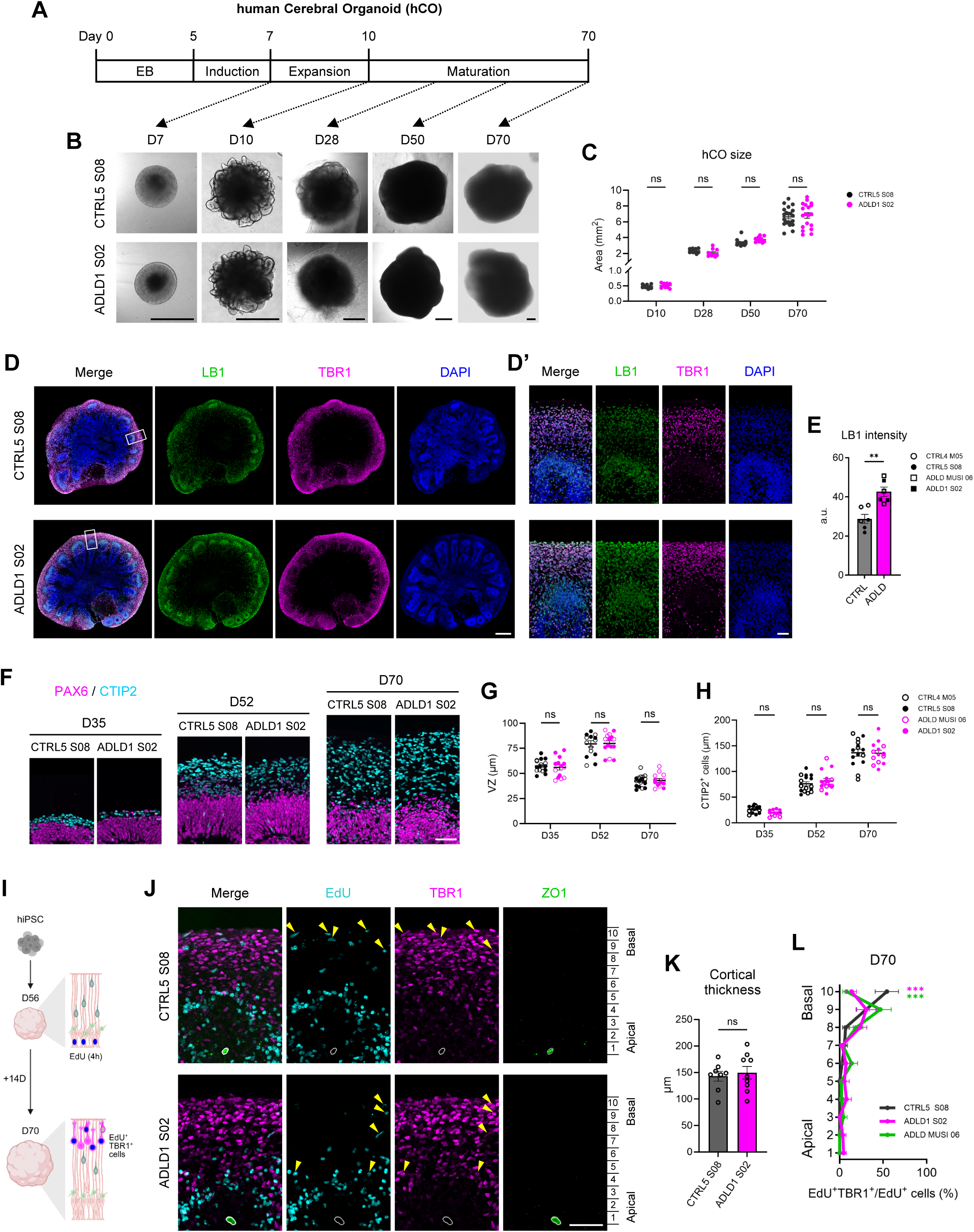
Impaired neuronal migration in brain organoids generated from ADLD patient-derived cells. **(A)** Timeline of hCO generation, including phases for embryoid body (EB) formation, neural induction, expansion, and maturation. **(B)** Representative bright-field images of live hCOs derived from CTRL and ADLD patient iPSCs at D7, D10, D28, D50, and D70. **(C)** Quantification of live hCO size (area) at the indicated time points. Counted organoids for CTRL5 S08 and ADLD1 S02 were 10 and 9 (D10); 10 and 10 (D28); 13 and 11 (D50); 22 and 19 (D70), respectively. Statistical analysis was performed using two-way ANOVA followed by Šidák’s multiple comparisons test. Error bars represent the mean ± SEM. **(D)** At D70, hCOs derived from CTRL and ADLD patient iPSCs were stained with LB1 (green) and TBR1 (magenta) antibodies, and counterstained with DAPI (blue). (D’) Enlarged views of the boxed regions in (D). **(E)** Quantification of LB1 protein intensity per organoid section (n = 2 individuals per group, 3 organoids analyzed per individual). Statistical analysis was performed using Student’s *t*-tests. Error bars represent the mean ± SEM. \*\**P* < 0.01. **(F)** Representative images of hCOs derived from CTRL and ADLD patient iPSCs, stained with PAX6 (magenta) and CTIP2 (cyan) antibodies at D35, D52, and D70. **(G)** Quantification of VZ thickness based on the PAX6+ region in (F) at D35, D52, and D70 (n = 2 individuals per group, 2–4 organoids analyzed per individual). Counted rosettes for CTRL and ADLD were 17 and 14 (D35); 15 and 15 (D52); 17 and 13 (D70). Statistical analysis was performed using two-way ANOVA followed by Šidák’s multiple comparisons test. Error bars represent the mean ± SEM. **(H)** Quantification of CTIP2+-layer thickness based on staining in (F) at D35, D52, and D70 (n = 2 individuals per group, 2–4 organoids analyzed per individual). Counted rosettes for CTRL and ADLD were 14 and 12 (D35); 16 and 16 (D52); 15 and 14 (D70). Statistical analysis was performed using two-way ANOVA followed by Šidák’s multiple comparisons test. Error bars represent the mean ± SEM. **(I)** Schematic of the EdU pulse-labeling to hCOs at D56 for 4 h, followed by fixation at D70. **(J)** Representative images of hCOs generated from CTRL and ADLD patient-derived iPSCs, stained with TBR1 (magenta) and ZO1 (green) antibodies along with EdU (cyan) detection at D70. The ZO1+ ventricular surface is outlined in white. Yellow arrowheads indicate EdU+TBR1+ neurons. **(K)** Quantification of cortical thickness measured from the ZO1+ ventricular surface to the surface of the hCOs. Statistical analysis was performed using Student’s *t*-tests. Error bars represent the mean ± SEM. **(L)** Quantification of the laminar distribution of EdU+TBR1+ neurons across bins 1 to 10, in which the ZO1 signal to the surface of the hCOs was divided into ten equal bins (n = 3 organoids per group). Counted cells for CTRL5 S08, ADLD1 S02, and ADLD MUSI 06 were 30, 33, and 44, respectively. Statistical analysis was performed using two-way ANOVA followed by Šidák’s multiple comparisons test. Error bars represent the mean ± SEM. \*\*\**P* < 0.001. Scale bars: 500 µm (B, D); 50 µm (D’, F, and J).

To assess neuronal migration, we conducted a birth dating assay in the hCOs using EdU, which is incorporated during the S-phase of mitotic NPCs (*45*). EdU was added to the culture medium for 4 hours to label the NPCs at D56, and the hCOs were fixed 14 days later (Fig. 8I). Subsequently, sections of the hCOs were immunostained with TBR1, a transcription factor expressed in newly born neurons (*46*). We quantified the distance of EdU-labeled TBR1^+^ neurons (EdU□TBR1□) from the ventricular surface, marked by ZO1-stained signals (Fig. 8J). Although the overall thickness from the ventricle to the surface of the hCOs did not differ between the two groups, we observed a significant shift in the spatial distribution of EdU□TBR1□ neurons (Fig. 8J–L). In the control hCOs, a greater number of newborn neurons reached the surface, whereas in the ADLD hCOs, they remained closer to the ventricle. Overall, our results indicate that the elevation of LB1 hinders the migration of human neurons within the confined environment of brain organoids.

## Discussion

### LB1 as a physical regulator in neuronal migration

*In vivo*, cell migration is driven by a dynamic interplay between cell-intrinsic machineries and the non-cell-autonomous microenvironment. Radial neuronal migration during cortical development is distinct from the migration of other cell types, such as cancer and immune cells (*13, 14*), in four key aspects: (i) the points of origin and destination are predetermined; (ii) neuronal migration must be completed within a defined developmental timeframe; (iii) neurons transform from a multipolar to a bipolar morphology to adhere to radial glial fibers before initiating locomotion; and (iv) they subsequently navigate through exceptionally narrow spaces occupied by the ECM, axons, and neighboring cells.

As with other cell types, successful neuronal migration critically depends on the translocation of the nucleus, the largest and stiffest organelle (*8*). Newly generated neurons express minimal levels of Lamin□A, the canonical regulator of nuclear stiffness (*20, 21*); however, they do express members of the Lamin□B family, particularly LB1 (Fig. 1L and S1). LB1 has been reported to maintain nuclear architecture in mouse embryonic fibroblasts (MEFs) derived from *Lmnb1*^Δ/Δ^ mice (*26*). In this study, we aimed to investigate whether LB1 serves as a key determinant of nuclear mechanics that limits nucleokinesis during passage through narrow intercellular spaces in the developing cortex.

To evaluate this hypothesis, we focused on the CP, where earlier-born neurons, after completing their migrations, create physical barriers that obstruct the radial migration of bipolar neurons. Three-dimensional measurements of all nuclei in the E18.5 mouse somatosensory cortex revealed that the mean distance from each nucleus to its three nearest neighbors was ∼7.27□µm, averaged across the lower, middle, and upper CP. This distance is smaller than the mean nuclear diameter (∼8.32□µm) in the same regions (Fig.□S5). Consequently, neurons must migrate through interstitial spaces narrower than their own nuclear size. These observations underscore the concept that nuclear deformability is a vital determinant of migratory efficiency in such confined environments.

To test this assertion, we manipulated the abundance of LB1 in a mosaic manner in the developing mouse cortex. LB1-OE neurons exhibited delayed migration, resulting in a subset of neurons arrested either beneath or below L2/3 of the CP. This phenomenon may arise from the intercellular spacing becoming narrower as it approaches the upper CP (Fig. 1B and S5C). At P0, LB1□enriched nuclei displayed a more spherical shape, correlating with the levels of LB1 protein (Fig.□2H). Moreover, AFM measurements revealed a corresponding increase in nuclear stiffness (Fig.□3D). Given that LB1 enrichment increases nuclear stiffness, we subsequently investigated how variations in stiffness impact nuclear deformability during migration. We built a computational model in which nuclear membrane stiffness was the variable, allowing us to assess migratory success over time (Fig. 4). The simulation revealed that when nuclear membrane stiffness increased, nuclear deformation was nearly eliminated, resulting in migration failure. Notably, the model predicted that nuclear elongation peaks just prior to the maximum increase in cellular velocity, followed by a subsequent relaxation (Fig. 4E). *In vivo* time-lapse imaging further validated this sequence; however, nuclear deformation persisted for several additional minutes, possibly due to the densely packed arrangement of neurons in the CP, which restricts complete relaxation (Fig.□5E and S6). This phenomenon is distinct from *in vitro* reports of “nuclear pistoning” in cancer cells migrating through microchannels (*47, 48*), emphasizing the dominant influence of the *in vivo* microenvironment.

In conclusion, these findings illustrate that successful radial migration requires significant nuclear deformation to navigate the narrow intercellular spaces within the CP. Notably, nuclear deformation is causally linked to cellular translocation by facilitating an increase in migration speed. Excess LB1 stiffens the nucleus, thereby inhibiting the process. This indicates that LB1-dependent nuclear mechanics is a principal rate-limiting factor in radial neuronal migration. While actomyosin contractility and cytoskeletal remodeling within the cell (*9, 49*), as well as cell-to-cell adhesion dynamics (*50, 51*), are important molecular functions for driving migration, our findings highlight nuclear deformation as a key physical constraint on neuronal movement within spatially restricted environments.

### Functions of LB1 in neuronal maturation

B□type Lamins are recognized as significant architects of chromatin organization and transcriptional regulation (*19*). Despite this, no significant changes in gene expression were detected in LB1-OE migrating neurons (Fig. 1L and M). The result implies that, during radial migration, LB1 primarily influences neuronal behavior via biophysical mechanisms rather than transcriptional pathways. At P14, when neuronal maturation is ongoing, LB1-OE neurons located in L2/3 exhibited electrical properties comparable to those of CTRL neurons. In contrast, ectopically positioned LB1-OE neurons showed significant deficits in dendritic elaboration and electrophysiological immaturity, even though they maintained the upper□layer molecular identity (Fig. 1E, G, and 6).

Previous studies on neocortical ectopia have demonstrated that mispositioned neurons are capable of receiving synaptic inputs from surrounding layers; however, they exhibit altered input–output patterns and restricted axonal projections, which disrupt the integration of laminar-specific circuit (*52*). Although LB1-OE in primary cultured neurons has been reported to reduce axonal outgrowth (*37*) and to mislocalize 14qD2L, a genomic locus containing genes critical for neuronal maturation (*53*), there is currently no direct evidence that LB1-OE cell-autonomously suppresses neuronal electrical activity. These findings suggest that the immaturity observed in ectopic LB1-OE neurons is primarily due to their inability to receive the appropriate laminar-specific cues necessary for post-migratory maturation.

At P14, LB1□KD neurons exhibited significantly reduced dendritic complexity, characterized by minimal branching and relatively short apical dendrites that did not fully extend (Fig. S4). This phenotype aligns with previous findings that LB1 plays a critical role in the later stages of neuronal maturation. For example, *Lmnb1* mutant mice display a thinner CP accompanied by severe laminar disorganization (*24, 26, 54*). *In vitro* studies have also demonstrated that LB1 KD impairs dendritogenesis, leading to reduced dendritic length and complexity (*37*). Combined with observations from LB1-OE neurons, these results underscore that both excess and deficiency of LB1 can disrupt neuronal morphology and function, thereby necessitating a precise balance of LB1 expression for proper neuronal maturation.

### Excessive LB1 in brain disorders

Mutations or dysregulation of Lamins are associated with various neurological disorders categorized as leukodystrophies (*55*). A prominent example is ADLD, primarily characterized by the duplication of the *LMNB1* gene (*27*) or deletions upstream of this gene (*28, 56*). These genetic alterations lead to elevated levels of LB1 (*56–58*). Clinically, ADLD typically manifests in mid-adulthood, with initial symptoms such as autonomic dysfunction, followed by progressive motor and cognitive decline (*59–61*). Although the precise mechanisms underlying the onset of the disease remain largely unclear, existing studies indicate that LB1-OE disrupts myelin maintenance, driving progressive vacuolar degeneration of white matter tracts. This degeneration is often accompanied by reactive astrocytic gliosis and secondary axonal loss, which are pathological hallmarks underlying the gradual but relentless clinical progression of ADLD (*29*).

Although ADLD has primarily been characterized as a glial-driven white matter disorder, previous reports suggest that LB1-OE may also adversely affect neuronal development during brain maturation (*37, 54, 62*). In our research utilizing cerebral organoid models derived from iPSCs of ADLD patients, we observed a delay in the radial migration of projection neurons. To date, the neuronal aspects of ADLD have received limited attention, largely overshadowed by its predominant glial pathology. One possible hypothesis is that mild laminar disorganization resulting from the subtle displacement of projection neurons may remain functionally silent during early life. Indeed, our patch-clamp analysis of mouse brains at P14/15 revealed that when LB1-OE neurons were substantially mispositioned outside of L2/3, either in the CP or the IZ, they exhibited functional abnormalities. However, LB1-OE neurons showed comparable electrophysiological properties to their controls as long as they were positioned within L2/3 (Fig. 6). As cortical circuitry undergoes continuous myelination and structural remodeling throughout the lifespan (*63, 64*), subsequent myelin degeneration could unmask these latent architectural defects. The cumulative impact of disrupted laminar organization and white matter loss may exacerbate network dysfunction, potentially contributing to the clinical onset and progression of ADLD. Moreover, we acknowledge that variations in nuclear stiffness may also influence glial migration within confined tissue environments, similar to neuronal migration. Such alterations could lead to glial misplacement and subsequent brain dysfunction in adulthood. It is imperative to conduct further research focusing on the pathological aspects of ADLD in relation to elevated LB1 levels, employing animal models and brain organoids to comprehensively address this critical issue.

## Materials and Methods

### Plasmids

The following plasmids were constructed in previous studies: pCAGGS (*65*); pCAG-EGFP-N1, pCAG-EGFP-C1, pCAG-H2B-TagRFP (*66*); pCXLE-*hOCT3/4*-*shp53*-F (Addgene, Cat# 27077); pCXLE-hSK (Addgene, Cat# 27078); pCXLE-hUL (Addgene, Cat# 27080); pCXWB-EBNA1 (Addgene, Cat#37624); and pCXLE-EGFP (Addgene, Cat# 27082). The human *LMNB1* cDNA (RefSeq: *NM_005573*) was amplified from a human cDNA library and subsequently cloned into pCAG-EGFP-N1 by replacing the EGFP sequence using XhoI and NotI restriction sites. The mouse *Lmnb1* cDNA (RefSeq: *NM_010721*) was amplified from pLV V5-Lmnb1 (Addgene, Cat# 175108) and inserted into pCAG-EGFP-C1 using NheI and SalI sites, replacing the EGFP sequence. For knockdown experiments, shRNA constructs targeting mouse *Lmnb1* were designed and cloned into pSilencer 3.0-H1 (NovoPro, Cat# V012700). Detailed information about these plasmids is provided in Table 1.

**Table 1.**
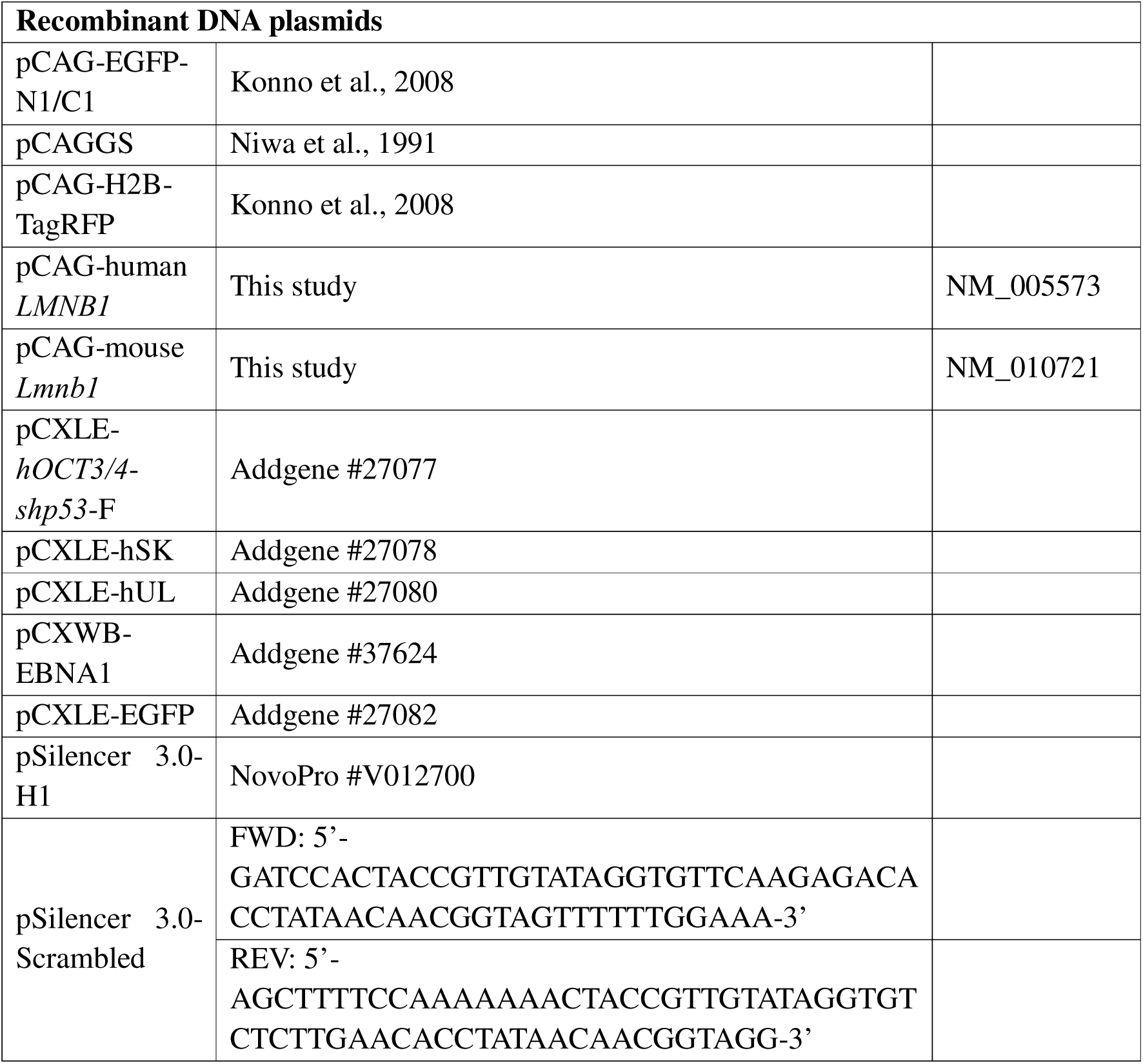
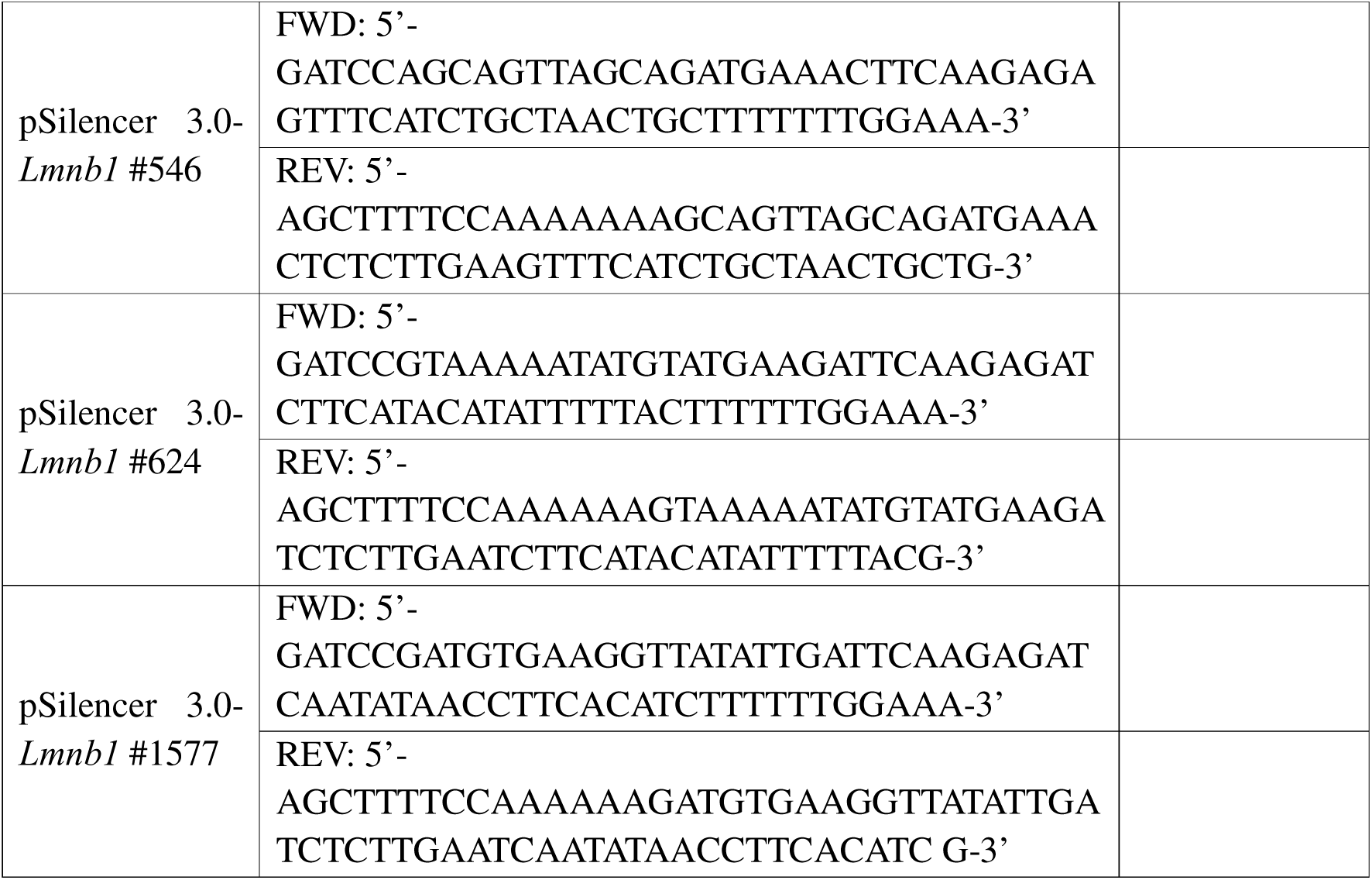
Plasmids.

### *In utero* electroporation (IUE)

IUE was performed at embryonic day (E) 14.5 or E15.5 as previously described (*67, 68*). The final plasmid solution contained 0.5 µg/µl of CAG-driven fluorescent protein expression vectors (pCAG-EGFP and pCAG-H2B-TagRFP) and 1 µg/µl of desired expression vectors (pCAGGS, pCAG-human *LMNB1*, and pCAG-mouse *Lmnb1*) with 0.01% fast green solution (Sigma-Aldrich, Cat# F7252). For E15.5 electroporation, electric pulses (35□V, 50□ms, 4 pulses) were applied using an electroporator (Nepa Gene, Cat#CUY21) and 5-mm forceps-type electrodes (Nepa Gene, Cat# CUY650P5). For E14.5 electroporation, 34□V pulses were used. After repositioning the embryos, the abdominal wall and skin were closed with sutures.

All animal experiments were approved by the Institutional Animal Care and Use Committee of the Korea Brain Research Institute (IACUC-22-00039-M3) and conducted in accordance with institutional guidelines.

### Mouse tissue preparation

Collected brains were fixed overnight at 4□°C in a solution of 4% paraformaldehyde (PFA) in 0.1□M phosphate buffer (PB). After sequential cryoprotection in 20% and 30% sucrose solutions in 0.1□M PB, the brains were embedded in Optimal Cutting Temperature (O.C.T.) compound (Sakura Finetek, Cat# 4583). Using a cryostat (Leica CM1860), the brains in the block were sectioned at 20–30 µm thickness. The sections were stored at −20□°C until immunostaining.

### Human tissue preparation

The fetal human brain was freshly frozen approximately 1.5 h after death. Sections were prepared by the Korea Brain Bank Network (KBBN). The sample was confirmed as a normal brain from a monozygotic, diamniotic monochorionic twin, based on the presence of fused placentas with visible superficial vascular anastomoses. Frozen cerebral cortical sections were transported on dry ice to the laboratory. After fixation in 4% PFA in PBS for 1 h, the tissues were immunostained with anti-Lamin B1 and anti-NeuN antibodies as described in the immunohistochemistry section.

All procedures involving human fetal tissue were approved by the Institutional Review Boards (IRB) of KBRI (KBRI-202111-BR-001-01) and KBBN (KBBN-00-DD01-21007) and were conducted in accordance with institutional and ethical guidelines.

### Immunohistochemistry (IHC)

IHC was performed on tissue sections and cultured cells with minor modifications to a previously described protocol (*69*). Frozen sections of mouse and human brain were brought to room temperature (RT) before undergoing antigen retrieval in HistoVT One solution (Nacalai Tesque, Cat# 06380-06) at either 70□°C or 90□°C for 20 min in a water bath. The sections were then washed twice with PBS, permeabilized with 0.05% Triton X-100 (Sigma-Aldrich, Cat# T8787) in PBS (PBSTx) for 10 min, and blocked with 2% bovine serum albumin (BSA; Sigma-Aldrich, Cat# A9418) in 0.05% PBSTx for 1 h at RT. For cell staining, the cells were fixed with 4% PFA for 10 min at RT, washed four times with PBS, and blocked with 2% BSA in 0.05% Triton X-100 in PBSTx for 1 h. All samples were incubated with primary antibodies overnight at 4□°C, washed three times with 0.05% Triton X-100 in PBSTx, and incubated with fluorescence-conjugated secondary antibodies and DAPI (Sigma, Cat# D9542) for 1 h at RT. Finally, the samples were mounted using ProLong Aqueous Mounting Medium (Thermo Fisher Scientific, Cat# P36930). Details regarding the antibodies are listed in Table 2. Fluorescence images were acquired using confocal microscopes (Andor Technology Dragonfly 502W; Leica TCS SP8 and STELLARIS 8).

**Table 2.** Primary and Secondary Antibodies.

| Primary antibodies |  |  |  |
| --- | --- | --- | --- |
| Lamin B1 | Rb | Abcam | ab16048; RRID: AB_443298 |
| Lamin B1 | Ms | Santa Cruz | SC-56144; RRID: AB_784272 |
| NeuN | Ms | Millipore | MAB377; RRID: AB_2298772 |
| CTIP2<br>(BCL11B) | Rat | Abcam | ab18465; RRID: AB_2064130 |
| SATB2 | Ms | Abcam | ab51502; RRID: AB_882455 |
| CUX1 | Rb | Proteintech | 11733-1-AP; RRID: AB_2086995 |
| PAX6 | Rb | MBL | PD022; RRID: AB_1520876 |
| TUBB3 | Ms | BioLegend | 801201; RRID: AB_2313773 |
| TBR1 | Rb | Abcam | ab31940; RRID: AB_2200219 |
| ZO-1 | Ms | Invitrogen | 33-9100; RRID: AB_2533147 |
| OCT4<br>(POU5F1) | Rb | Abcam | ab19857; RRID: AB_445175 |
| NANOG | Rb | MBL | PM058 |
| SOX2 | Rb | Millipore | AB5603; RRID: AB_2286686 |
| TRA-1-81 | Ms | Millipore | MAB4381; RRID: AB_177638 |
| SSEA-4 | Ms | Santa Cruz | sc-21704; RRID: AB_628289 |
| **Secondary antibodies** |  |  |  |
| Goat anti-Rat IgG (H+L), Alexa Fluor 555 |  | Invitrogen | A21434 |
| Goat anti-Rabbit IgG (H+L), Alexa Fluor 647 |  | Invitrogen | A21245 |
| Goat anti-Rabbit IgG (H+L), Alexa Fluor 568 |  | Invitrogen | A11036 |
| Goat anti-Rabbit IgG (H+L), Alexa Fluor 555 |  | Invitrogen | A21428 |
| Goat anti-Mouse IgG (H+L), Alexa Fluor 647 |  | Invitrogen | A21235 |
| Goat anti-Rabbit IgG (H+L), Alexa Fluor 488 |  | Invitrogen | A32731 |
| Goat anti-Mouse IgG (H+L), Alexa Fluor 555 |  | Invitrogen | A21422 |
| Goat anti-Mouse IgG (H+L), Alexa Fluor 488 |  | Invitrogen | A11001 |

### Primary culture

The cellular dissociation of EGFP^+^ dorsal cortical tissue and the subsequent culture conditions were performed with minor modifications to a previously described method (*39*). The dissociated cells were plated onto glass-bottom or plastic dishes pre-coated with 15□µg/mL poly-L-ornithine (Merck Millipore, Cat# P3655) and 1□µg/mL fibronectin (R&D Systems, Cat# 1030-FN). Cells were seeded at a density of 5.7 × 10□ cells/cm².

### FACS

Cortices were dissected from E18.5 mouse embryos 3 days after IUE at E15.5. After removal of the meninges, the tissues were placed in ice-cold DMEM/F-12 (Thermo Fisher Scientific, Cat# 11320033). The EGFP^+^ tissues were enzymatically dissociated using Accutase (Innovative Cell Technologies, Cat# AT-104) at 37°C for 10 min. Tissues were further dissociated by gentle mechanical trituration and centrifuged at 200 × g for 3 min. The cell pellet was resuspended in DMEM/F-12 and passed through a 40-μm cell strainer (Falcon, Cat# 352340), followed by centrifugation under the same conditions. The final cell pellet was resuspended in DMEM/F-12 supplemented with 5 mM EDTA (Thermo Fisher Scientific, Cat# 15575038) and 0.5% N-2 supplement (Thermo Fisher Scientific, Cat# 17502048). EGFP^+^ cells were then isolated using a FACSAria III cell sorter (BD Biosciences). The sorted cells were collected directly into DMEM/F-12 containing 0.5% BSA (Thermo Fisher Scientific, Cat# 15260037). Total RNA was extracted using the RNeasy Plus Mini Kit (Qiagen, Cat# 74136) following the manufacturer’s instructions. RNA integrity was assessed using the High Sensitivity RNA ScreenTape system (Agilent, Cat# 5067), and only samples with RNA integrity number (RIN) values 7.0 or higher were used for further analysis (mean RIN = 7.7). To ensure biological reproducibility, two independent IUE and FACS experiments were conducted. In the first experiment, 19 CTRL and 22 hLB1 embryos were obtained from 3 pregnant ICR mice. In the second experiment, 24 CTRL and 28 hLB1 embryos were obtained from 4 pregnant ICR mice.

### RNA-sequencing analysis

Library preparation for bulk RNA samples was conducted using the FLASH-seq method (*70*). Sequencing was performed on the Illumina NextSeq 2000 platform, generating paired-end reads of 36 bp. The sequenced reads were aligned to the mouse (UCSC-mm10) and human (UCSC-hg38) reference genomes using STAR version 2.7.1a with some options (--outFilterMultimapNmax 20 --alignSJoverhangMin 8 --alignSJDBoverhangMin 1 -- outFilterMismatchNmax 999 --alignIntronMin 20 --alignIntronMax 10000 -- alignMatesGapMax 1000000). Aligned reads per gene were quantified using featureCounts from the subread package version 2.0.1. Transcript abundances were normalized to transcripts per million (TPM). Differentially expressed genes (DEGs) were identified using DESeq2 version 1.42.1 in R. Genes with an adjusted P-value of less than 0.05 and a |log2 fold change| greater than 1 were considered significantly differentially expressed.

### Atomic force microscopy (AFM)

AFM measurements were conducted with slight modifications to a previously described protocol (*39*). We utilized an atomic force microscope (BioScope Resolve; Bruker), controlled with NanoScope 9.4 software and mounted on an inverted microscope (Nikon, ECLIPSE Ti2). A tipless silicon cantilever equipped with a 20-μm borosilicate bead (Novascan, Cat# PG20T) was employed. The spring constant of each cantilever was calibrated in air using the thermal noise method. Cantilevers with consistent spring constants (nominal: 0.03 N/m; actual: 0.04 N/m) were selected for the measurements, and a constant force of 5 nN was applied. Force–distance curves were obtained in contact mode. Both fluorescence and bright-field images were captured using a CMOS camera (Hamamatsu, ORCA-Flash4.0, Cat# C13440-20CU) to identify the cells being measured, with EGFP^+^ cells randomly selected for analysis. Young’s modulus was calculated by fitting the force–distance curves to the Hertz model using NanoScope Analysis 1.9 software (Bruker).

### Transwell migration assay

The transwell migration assay was conducted with slight modifications to a previously described protocol (*71*). Cells dissociated from the EGFP^+^ dorsal cortex were seeded onto transwell inserts with an 8-μm pore size (Corning, Cat# 3422), which had been pre-coated with 15□μg/mL poly-L-ornithine (Merck Millipore, Cat# P3655) and 1□µg/mL fibronectin (R&D Systems, Cat# 1030-FN). For the assay, 200 µl of medium consisting of Neurobasal (Thermo Fisher Scientific, Cat# 21103049) supplemented with 2% B-27 Supplement (Thermo Fisher Scientific, Cat# 17504044) and 1% penicillin–streptomycin (Thermo Fisher Scientific, Cat# 15140122), containing 5 × 10□ cells, was added to the upper chamber of each insert. Simultaneously, 600□µL of the same medium was added to the lower chamber. After 40 h of incubation, the cells were fixed with 4% PFA in PBS for 10 min. Each insert was then carefully removed, mounted on a glass-bottom dish, and imaged using a confocal microscope (Leica, TCS SP8).

### Cell arrest assay

The cell arrest assay was conducted with slight modifications to a previously described protocol (*15*). Cells dissociated from the EGFP^+^ dorsal cortex were resuspended in a medium consisting of Neurobasal (Thermo Fisher Scientific, Cat# 21103049) supplemented with 2% B-27 Supplement (Thermo Fisher Scientific, Cat# 17504044), 1% N-2 Supplement (Thermo Fisher Scientific, Cat# 17502048), 0.5% penicillin–streptomycin (Thermo Fisher Scientific, Cat# 15140122), 1% GlutaMAX (Thermo Fisher Scientific, Cat# 35050061), and 0.1□mM Trolox (Sigma-Aldrich, Cat# 238813) to achieve a final density of 1 × 10□ cells/ml. The cell suspension was then introduced into the microfluidic device at a flow rate of 1□µl/min using a syringe pump (New Era Pump Systems, Cat# NE-1000).

### Time-lapse imaging

Time-lapse imaging was conducted with slight modifications to previously described protocols (*72*). Briefly, IUE was performed at E15.5 in the cortex of ICR mice using pCAG-EGFP and the desired plasmids. Three days later, the brains were collected, embedded in 3% low-melting-point agarose (Duchefa Biochemie, Cat# L1204), and sectioned coronally at a thickness of 300□µm using a vibratome (Leica VT1200S). The brain slices were placed on Millicell-CM culture plate inserts (Merck Millipore, Cat# PICMORG50), positioned on glass-bottom dishes (MatTek, Cat# P35G-1.5-20-C). The slices were cultured in Neurobasal medium (Thermo Fisher Scientific, Cat# 21103049) supplemented with 2% B-27 Supplement (Thermo Fisher Scientific, Cat# 17504044), 1% N-2 Supplement (Thermo Fisher Scientific, Cat# 17502048), 0.5% penicillin–streptomycin (Thermo Fisher Scientific, Cat# 15140122), 1% GlutaMAX (Thermo Fisher Scientific, Cat# 35050061), and 0.1□mM Trolox (Sigma-Aldrich, Cat# 238813). Imaging was performed using a spinning disk confocal microscope (Andor Technology Dragonfly 502W) equipped with a temperature- and CO□-controlled chamber (37□°C, 5% CO□). Approximately 60–70 optical Z-sections were acquired every 5 min for over 24 h, and time-lapse movies were generated by projecting all Z-planes into a maximum intensity projection at each time point.

### Electrophysiological recordings

Acute brain slices were prepared from P14–15 mice. The mice were anesthetized using sodium pentobarbital (70 mg/kg), and the brain tissues were quickly extracted for slice preparation. The brains were immediately removed and immersed in a chilled cutting solution with the following composition (in mM): 110 choline chloride, 2.5 KCl, 25 NaHCO_3_, 1.25 NaH_2_PO_4_, 25 glucose, 0.5 CaCl_2_, 7 MgCl_2_·6H_2_O, 11.6 sodium L-ascorbate, and 3 pyruvic acid. Subsequently, 300-µm-thick coronal slices were prepared using a vibratome (Leica, VT1200S). The slices were then incubated for 30 min at 32 °C in artificial cerebrospinal fluid (ACSF) with the following composition (in mM): 119 NaCl, 2.5 KCl, 26 NaHCO_3_, 1.25 NaH_2_PO_4_, 20 glucose, 2 CaCl_2_, 1 MgSO_4_, 0.4 L-Ascorbic acid, and 2 pyruvic acid. During the slicing process, the solutions were saturated with carbogen (95% O2, 5% CO2) to maintain a final pH of 7.4.

Whole-cell patch-clamp recordings were conducted to the slices placed in a submerged recording chamber with a continuous flow of ACSF saturated with carbogen. Layer 2/3 neurons of the medial prefrontal cortex (mPFC) were visualized using an upright microscope (Olympus BX51WI) equipped with differential interference contrast (DIC) optics and a 40x water immersion objective (NA 0.8, Olympus). All electrophysiological recordings were conducted at 26 ± 2 °C, with fresh ACSF perfused at approximately 1.5 ml/min. The patch electrodes were pulled from borosilicate glass capillaries to achieve a resistance of 4 to 5 MΩ. The internal solution contained the following components (in mM): 138 potassium gluconate, 10 KCl, 10 HEPES, 10 Na2-phosphocreatine, 4 MgATP, 0.3 NaGTP, and 0.2 EGTA, adjusted to a pH of 7.25.

### Preparation of human dermal fibroblasts from patients and controls

Primary human fibroblasts isolated from skin biopsies of ADLD patients carrying *LMNB1* gene duplication and non-diseased donors were obtained from the IRCCS Istituto delle Scienze Neurologiche di Bologna, UOC NeuroMet, Italy. This study was approved by the AUSL Bologna Ethical Committee and informed consents were obtained from all participants.

Briefly, after collection, the samples were immediately submerged in complete DMEM supplemented with antibiotic-antimycotic solution (Thermo Fisher Scientific, Cat# 15240062). The samples were then cut into fragments using a sterile scalpel, and each fragment was placed in a marked Petri dish. The marked area was created by scratching the surface of the dish to enhance specimen attachment. Samples were incubated in a humidified incubator at 37□°C with 5% CO□. The culture medium was changed every 2–3 days. Cells were passaged using Accutase (Thermo Fisher Scientific, Cat# 00-4555-56) when they reached 80% confluence.

### Generation and maintenance of hiPSCs

Induced pluripotent stem cell (iPSC) generation was conducted with slight modifications to a previously described protocol (*73*). A Y4 episomal plasmid mixture was prepared, consisting of pCXLE-hOCT3/4-shp53-F (Addgene, Cat# 27077), pCXLE-hSK (Addgene, Cat# 27078), pCXLE-hUL (Addgene, Cat# 27080), pCXWB-EBNA1 (Addgene, Cat# 37624), and pCXLE-EGFP (Addgene, Cat# 27082). This mixture was then electroporated into HDFs using a Nepa21 electroporator (Nepa Gene) with the following settings: a poring pulse voltage of 200 V, a pulse duration of 5 ms, an interval of 50 ms, and two pulses. Following electroporation, the cells were plated onto either SNL feeder layers (Cell Biolabs, Cat# CBA-316) or MEF feeder layers (Cell Biolabs, Cat# CBA-312). The cells were maintained in an ES medium composed of DMEM/F12 (Thermo Fisher Scientific, Cat# 11320-033) supplemented with 4ng/ml of bFGF (Peprotech, Cat# 100-18), 20% KSR (Thermo Fisher Scientific, Cat# 10828028), 1% MEM-NEAAs (Thermo Fisher Scientific, Cat# 11140050), and 0.1% beta-mercaptoethanol (Thermo Fisher Scientific, Cat# 21985023) until iPSC colonies emerged. Once established, the iPSC colonies were transferred to a feeder-free culture system and subsequently maintained in mTeSR medium. Karyotype analysis was performed by Samkwang Medical Laboratory (Korea). All procedures involving human cells were approved by the IRB of KBRI (KBRI-202502-BR-001-01E) were conducted in accordance with institutional and ethical guidelines.

### Neural differentiation

Neural differentiation was carried out with slight modifications to previously described protocols (*45, 74*). To initiate the differentiation process (designated as neural differentiation day 0; ND0), the culture medium was replaced with neural induction medium consisting of DMEM/F12 + GlutaMAX (Thermo Fisher Scientific, Cat# 10565018) supplemented with 2% B-27 Plus (Thermo Fisher Scientific, Cat# A3582801), 1% N-2 Supplement (Thermo Fisher Scientific, Cat# 17502048), 0.5% penicillin–streptomycin (Thermo Fisher Scientific, Cat# 15140122), and 100 ng/mL recombinant human Noggin (PeproTech, Cat# 120-10C). This medium was changed every 2 days until ND16. From ND16 to ND24, the same medium was used without Noggin. From ND24 onward, the medium was switched to a neuronal maintenance medium composed of Neurobasal Plus (Thermo Fisher Scientific, Cat# A3582901) supplemented with 2% B-27 Supplement without retinoic acid (Thermo Fisher Scientific, Cat# 12587010), 1% GlutaMAX (Thermo Fisher Scientific, Cat# 35050061), and 0.5% penicillin–streptomycin. The medium was changed every 3–4 days until ND40.

### Quantitative real-time PCR (qPCR)

On a designated day during neural differentiation, total RNA was extracted using the RNeasy Plus Mini Kit (Qiagen, Cat# 74136), following the manufacturer’s instructions. Quantitative real-time PCR was performed with slight modifications to a previously described protocol (*75*). Briefly, equal amounts of cDNA were synthesized from the total RNA using the ReverTra Ace qPCR RT Kit (TOYOBO, Cat# FSQ-201). The qPCR was conducted using the FastStart Universal SYBR Green Master Mix (Roche, Cat# 0913850001) with the following cycling conditions: 95□°C for 10 min, followed by 40 cycles of 95□°C for 15 sec and 60□°C for 60 sec, performed on a LightCycler 480 II system (Roche). GAPDH was used as an internal control. Amplification specificity was confirmed by melt curve analysis. The primer sequences are listed in Table 3.

**Table 3.**
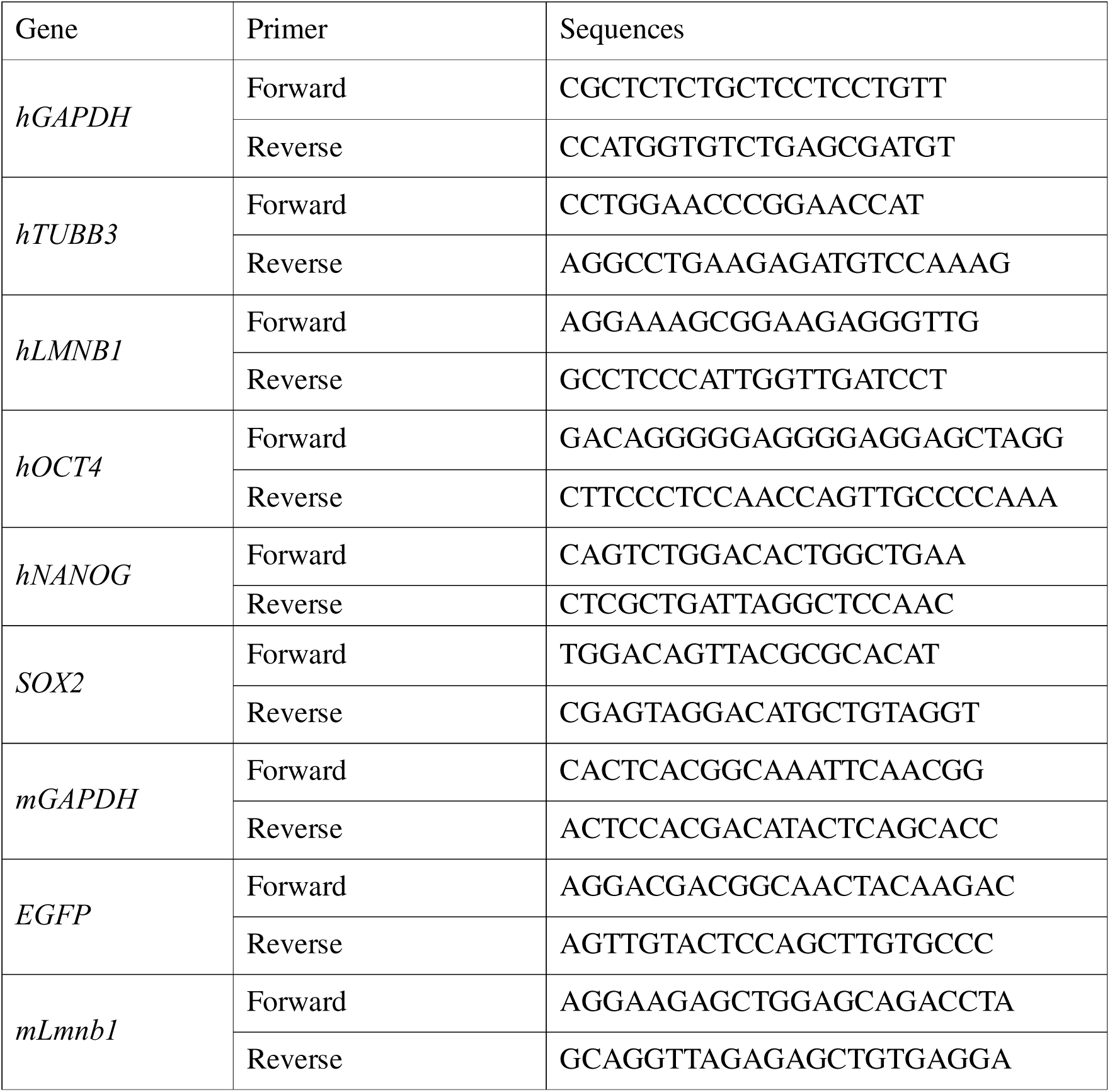
Primers.

### Cerebral organoid (CO) generation and EdU incorporation

COs were generated using the STEMdiff Cerebral Organoid Kit (Stemcell Technologies, Cat# 08570), following the manufacturer’s instructions. On Day 0, hiPSCs maintained in mTeSR medium (Stemcell Technologies, Cat# 05850) were dissociated using Accutase (Innovative Cell Technologies, Cat# AT-104) and seeded into ultra-low attachment 96-well plates (Corning, Cat# 7007) at a density of 2.0 × 10□ cells per well in embryoid body (EB) formation medium supplemented with 10 µM Y-27632 (FUJIFILM Wako, Cat# 036-24023). The medium was changed every 2 days without Y-27632. To induce cerebral organoid formation, one EB per well was transferred to a 24-well ultra-low attachment plate (Corning, Cat# 3473) containing induction medium. After 7 days, the organoids were embedded in Matrigel (Corning, Cat# 354230) and cultured in expansion medium. After 4 weeks, the medium was switched to neuron differentiation medium, and the organoids were further cultured for up to 70 days (*76*). For the cell migration assay, EdU (Thermo Fisher Scientific, Cat# C10340) was added to the culture medium on Day 56 and incubated for 4 h. Subsequently, the medium was replaced with fresh medium without EdU. Fixation was performed 14 days later to trace the EdU-labeled cells.

### Computational modeling

Neuronal migration in a densely packed environment was modeled as a cell migrating through immobile obstacles. The cell was modeled as a soft capsule containing a smaller, harder core inside, simulating a neuron with a nucleus. The cell and nucleus, and the cellular environment, were assumed to be filled with a Newtonian liquid of viscosity η. The deformability of the cell and nucleus was characterized by the membrane stiffness, or the surface shear elastic modulus G_s_. The cell and nucleus membranes were assumed to follow the strain energy function (*77*):

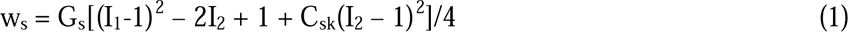

where I_1_ and I_2_ are the invariants of the surface right Cauchy-Green deformation tensor, and C_sk_ = 10^2^ is the area-dilation factor. The bending resistance (*78*) was also considered for the cell and nucleus membranes. The cell was initially spherical, with a diameter of d = 10 μm, and the nucleus was also spherical, with a diameter of 6 μm. The immobile obstacles were modeled as fixed rigid spheres with a diameter of 10 μm. The fluid mechanics was solved by the lattice Boltzmann method (*79*), the membrane mechanics was solved by the finite element method (*80*), and they were coupled by the immersed boundary method (*81*). For the numerical methods, see our previous papers (*82–84*). In the simulation, the cell movement was driven by body force. We varied the ratio of the nucleus membrane stiffness to the cell membrane stiffness RG_s_, and the spacing between the immobile obstacles δ. Displacement was normalized as x/d, and velocity was ηv/G_s_.

### Image analysis

Images were acquired using confocal microscopes, specifically the Dragonfly 502W (Andor Technology), TCS SP8, and STELLARIS 8 (Leica Microsystems). Z-stack images were captured using 10x, 20x, or 40x objectives, chosen according to experimental requirements. Cell position, nuclear morphology, marker positivity, cell processes, and LB1 intensity were analyzed using ImageJ 1.54p (Fiji; Wayne Rasband and contributors, National Institutes of Health, USA). For transwell assays, Aivia 13.1 (Leica) was used to determine the Z-axis center of mass of EGFP^+^ cells. Time-lapse image analysis was performed using a combination of Imaris 9.7 (Bitplane) and ImageJ 1.54p. The Y-positions, displacement lengths, and migration velocities of EGFP^+^ cells were measured using the *Spots* function in Imaris. Nuclear shape, estimated from the H2B-TagRFP signal, was analyzed in ImageJ 1.54p. Three-dimensional reconstruction of DAPI-stained images in Imaris 9.7 using *Spots* and *Surfaces* functions from 110 optical sections acquired at 1.0-µm z-steps.

### Statistical analysis

Statistical analyses and graph visualization were performed using GraphPad Prism version 10.5. Statistical significance for two-group comparisons was assessed using two-tailed unpaired Student’s *t*-tests. For multiple-group comparisons, one-way or two-way analysis of variance (ANOVA) was followed by Dunnett’s multiple comparisons test when comparing each experimental group with the control, by Tukey’s multiple comparisons test for all pairwise comparisons, or by Šidák’s multiple comparisons test for planned pairwise contrasts. A *P*-value < 0.05 was considered statistically significant, whereas values > 0.05 were considered not significant (ns). Maximum *P*-value indicated here is < 0.001. The specific experimental conditions and statistical tests are described in the corresponding figure legends.

## Acknowledgments

The brain research resources (human brain tissue) were provided by Korea Brain Bank Network (Asan Medical Center Brain Bank) operated through National Brain Bank Project(**25-BR-09-01**) and Korean Brain cluster promotion project(**RS-**2021**-NR057633**), both funded by the MSIT. We would like to express our sincerest gratitude to Eun-Jae Lee, Soo Jeong Nam (Asan hospital) for providing human brain sections; Prof. Kazunori Nakajima□ (Keio University), Prof. Ken□ichiro Kubo, and Dr. Ayako Kitazawa (The Jikei University School of Medicine) for IUE and time-lapse guidance; Gwanghyun□Park for iPSC setup; Dr. Hiroki□Sugishita and Ayako□Watanabe (The University□of□Tokyo) for FLASH□seq; Vittoria□Piancaldini and Foteini□Dionysia□Koufi (University of Bologna) for hiPSC characterization; Youngjae□Ryu (KBRI) for confocal imaging; Soojin□Jo (DGIST) for FACS assistance.

## Author contributions

M.S. and Y.K. designed the study. M.S. performed experiments. S.I. and Y.I performed computational modeling. J.Y. and J.R. performed patch-clamp assay. M.I. performed AFM studies. G.J. performed FLASH-seq data analysis. P.C., E.G., I.C., G. R. and S.R. prepared CTRL and ADLD patient-derived cells. D.Y. supervised cell arrest assay. M.S. and Y.K. wrote the manuscript with input from all authors. Y.K. supervised the project.

