## Supplemental Figures for "Lamin B1 physically regulates neuronal migration by modulating nuclear deformability in the developing cortex"

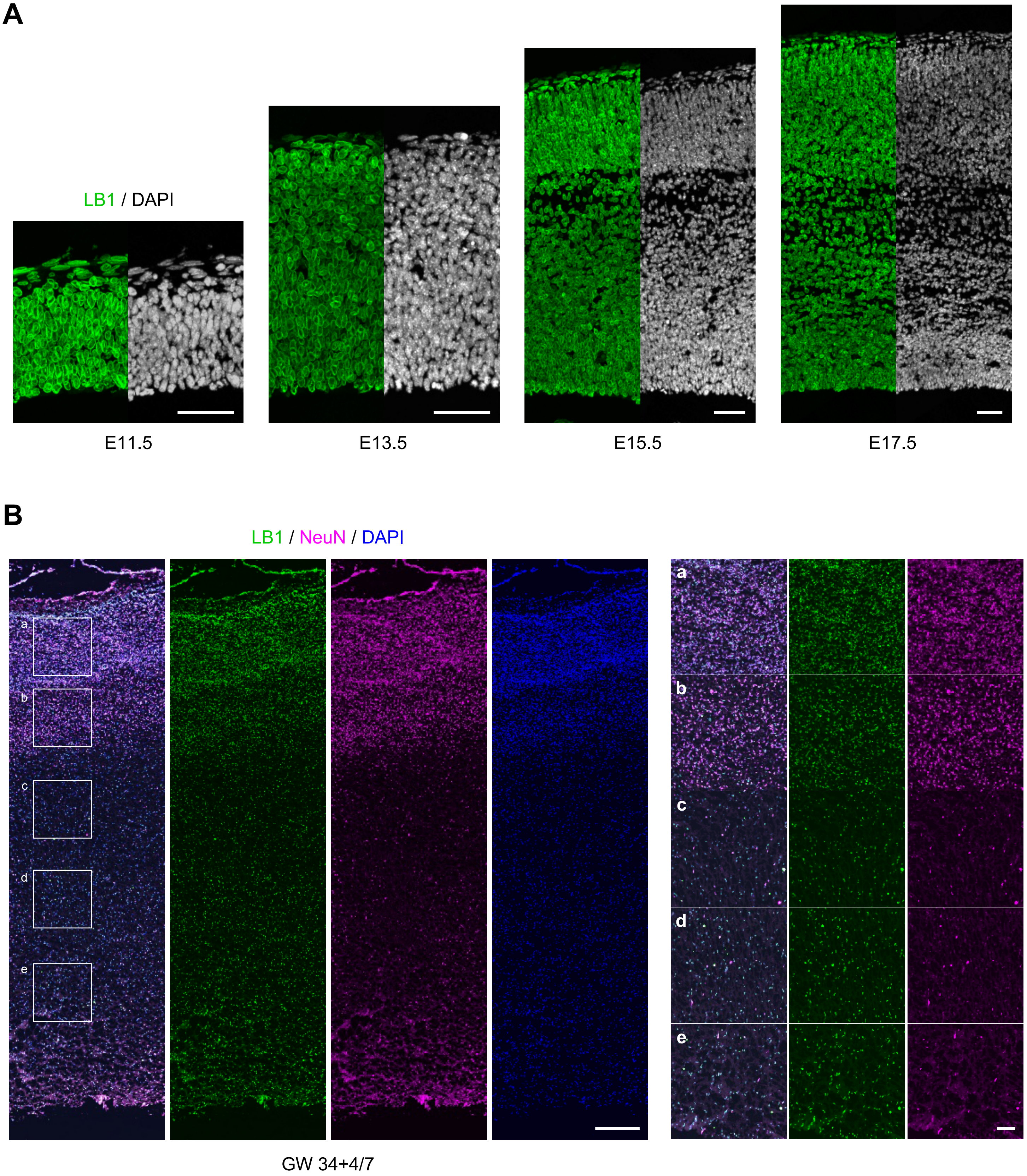


**Fig. S1. Expression patterns of LB1 protein during mammalian cortical development.**
(**A**) Coronal brain sections from the mouse neocortex at E11.5, E13.5, E15.5, and E17.5 were stained with LB1 antibody (green) and counterstained with DAPI (blue).
(**B**) A coronal brain sections from human neocortex at GW34+4/7 was stained with LB1 (green) and NeuN (magenta) antibodies, and counterstained with DAPI (blue). The white boxes indicate regions shown at higher magnification.
Scale bars: 50 μm (A); 500 μm (B); 100 μm (B').

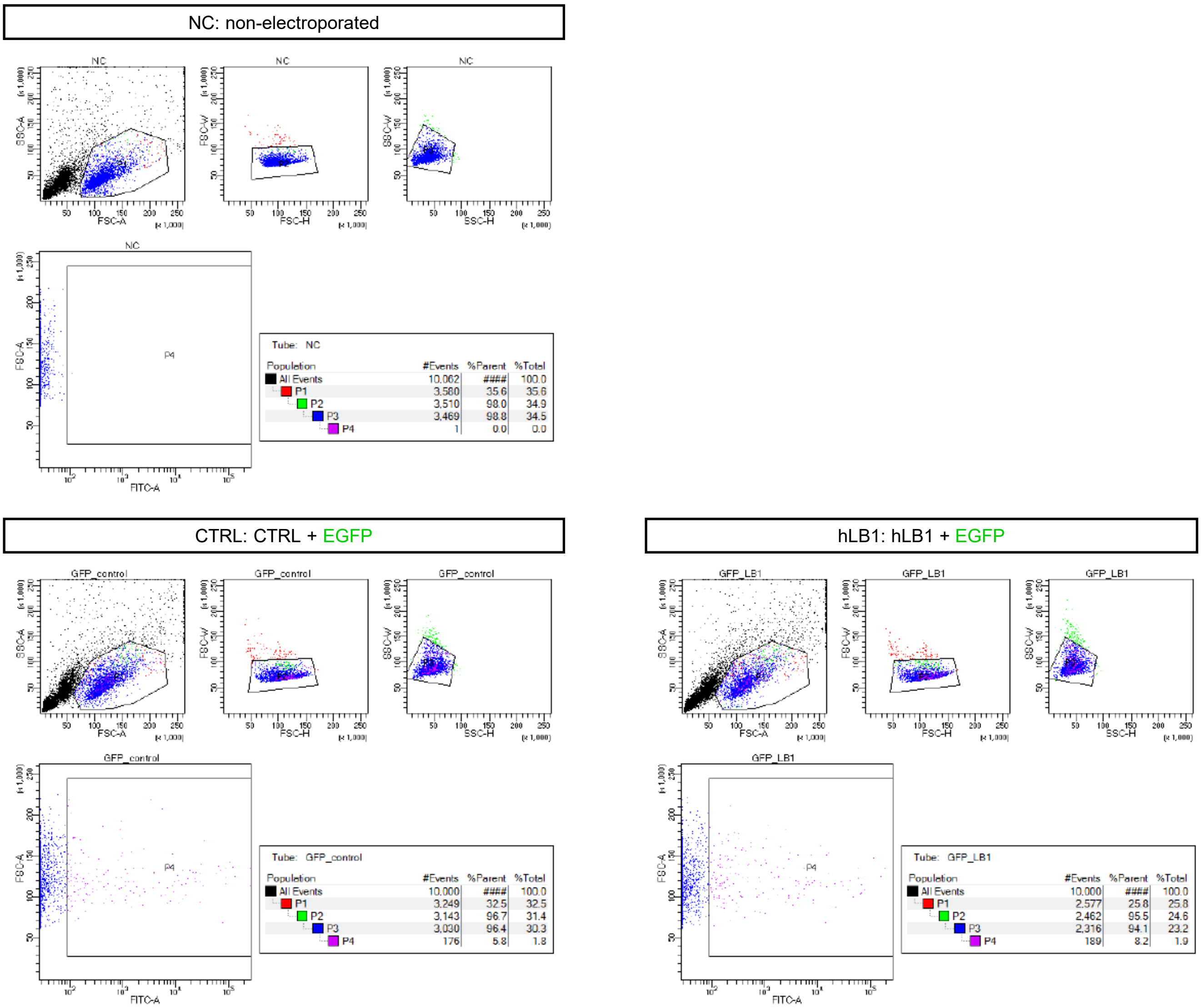


**Fig. S2. FACS analysis of EGFP+ cell sorting following IUE.**

Examples of FACS plots of non-electroporated (NC), CTRL, and hLB1-OE cells obtained from the E18.5 cortex after IUE at E15.5, related to Fig. 1I–M.

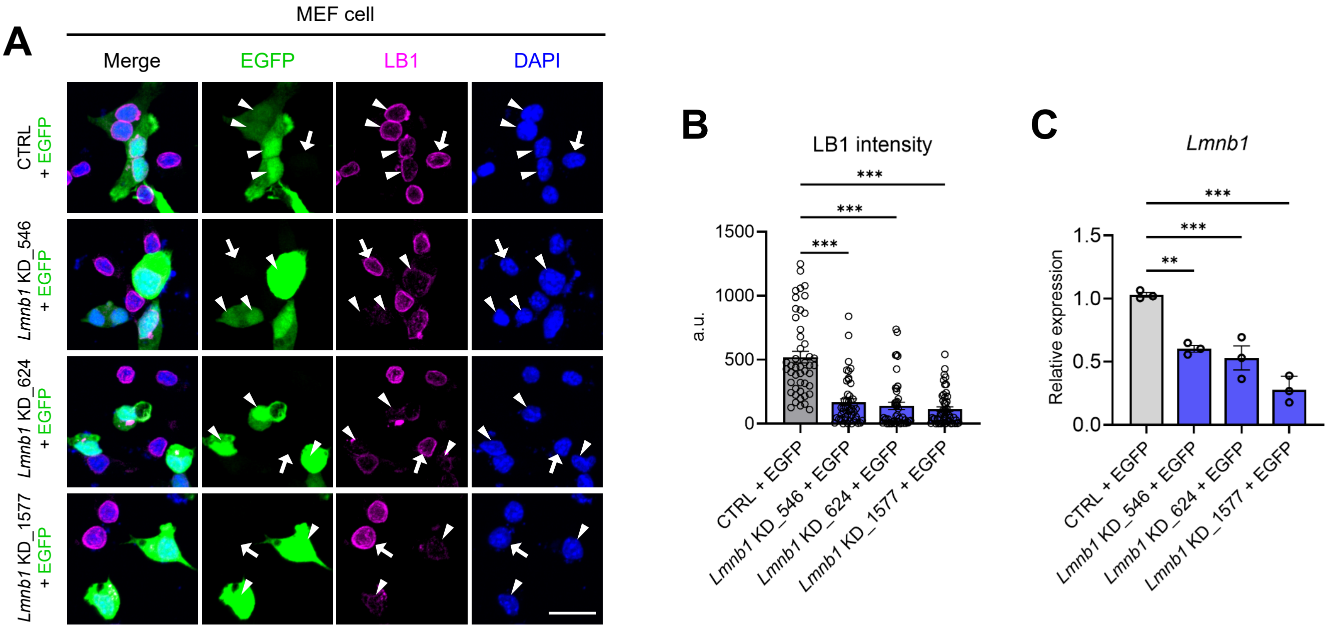


**Fig. S3. Knockdown of *Lmnb1* reduces endogenous LB1 protein and mRNA levels in MEF cells.**

(**A**) MEF cells were electroporated with scrambled control (CTRL) or Lmnb1-targeting shRNA (Lmnb1 KD) plasmids with pCAG-EGFP. The cells were fixed after 48 h and stained with LB1 antibody (magenta). Arrowheads, EGFP+ cells; arrows, EGFP- cells.

(**B**) Quantification of LB1 protein intensity in EGFP+ cells in (A). Counted cells for CTRL, Lmnb1 KD_546, Lmnb1 KD_624, and Lmnb1 KD_1577 were 47, 47, 46, and 51, respectively.

(**C**) Quantification of relative mRNA expression levels of Lmnb1 measured by qPCR at 48 h after electroporation. Each dot represents an independent experiment.

Based on these results, Lmnb1 KD_1577 was selected for subsequent Lmnb1 KD experiments in Fig. 2M–V.

Statistical analysis was performed using one-way ANOVA followed by Tukey's multiple comparison test. Error bars represent the mean ± SEM. $^{**}P < 0.01$, $^{***}P < 0.001$.

Scale bar: 20 μm (A).

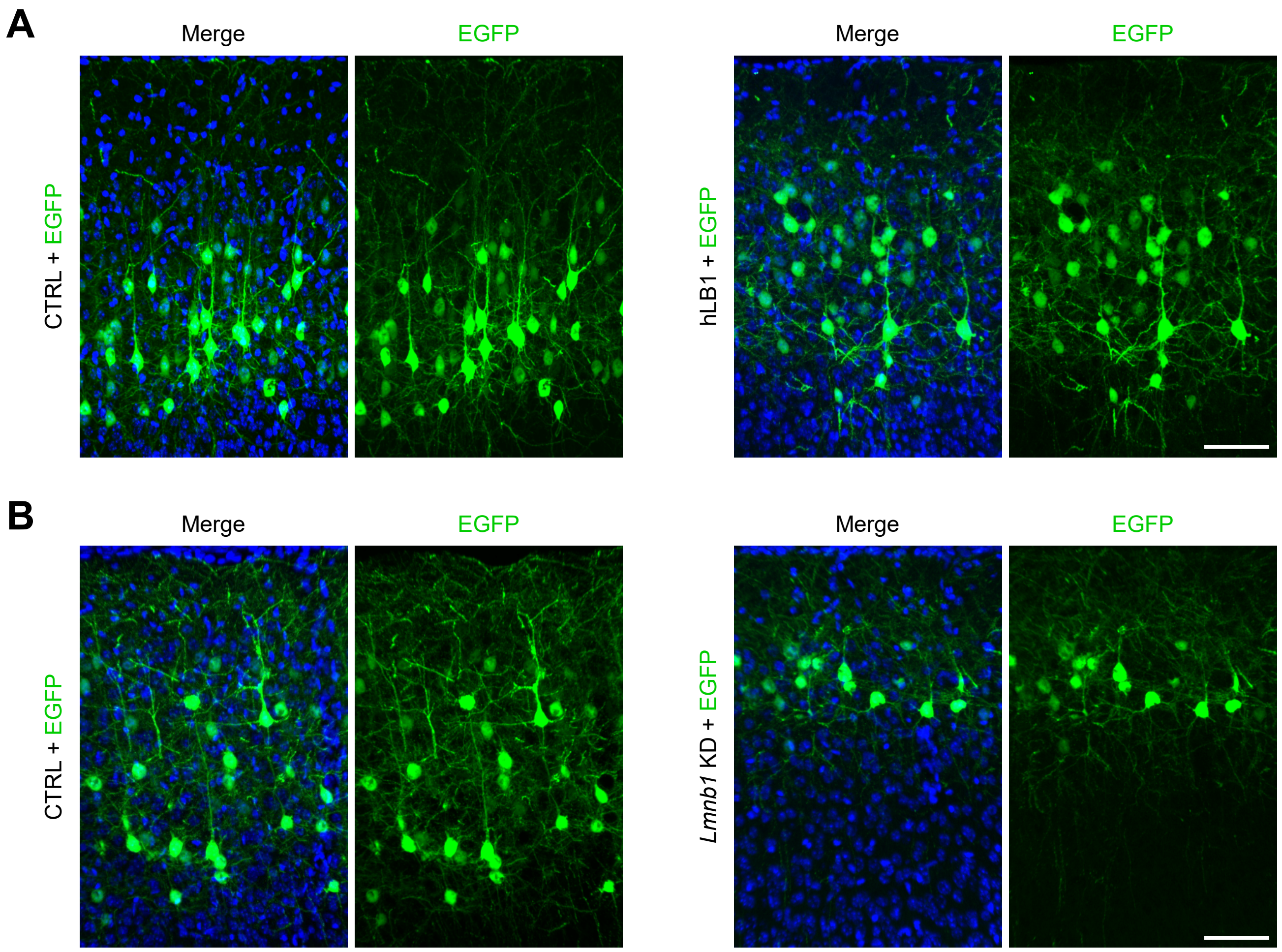


**Fig. S4. Neuronal morphology of CTRL, hLB1-OE, and *Lmnb1* KD groups.**
(**A**) The neocortex of E15.5 mice was electroporated with CTRL or hLB1 plasmids with pCAG-EGFP, then fixed at P14. Coronal brain sections were counterstained with DAPI.
(**B**) The neocortex of E15.5 mice was electroporated with CTRL or Lmnb1 KD plasmids with pCAG-EGFP, then fixed at P14. Coronal brain sections were counterstained with DAPI.
Scale bar: 80 μm (A, B).

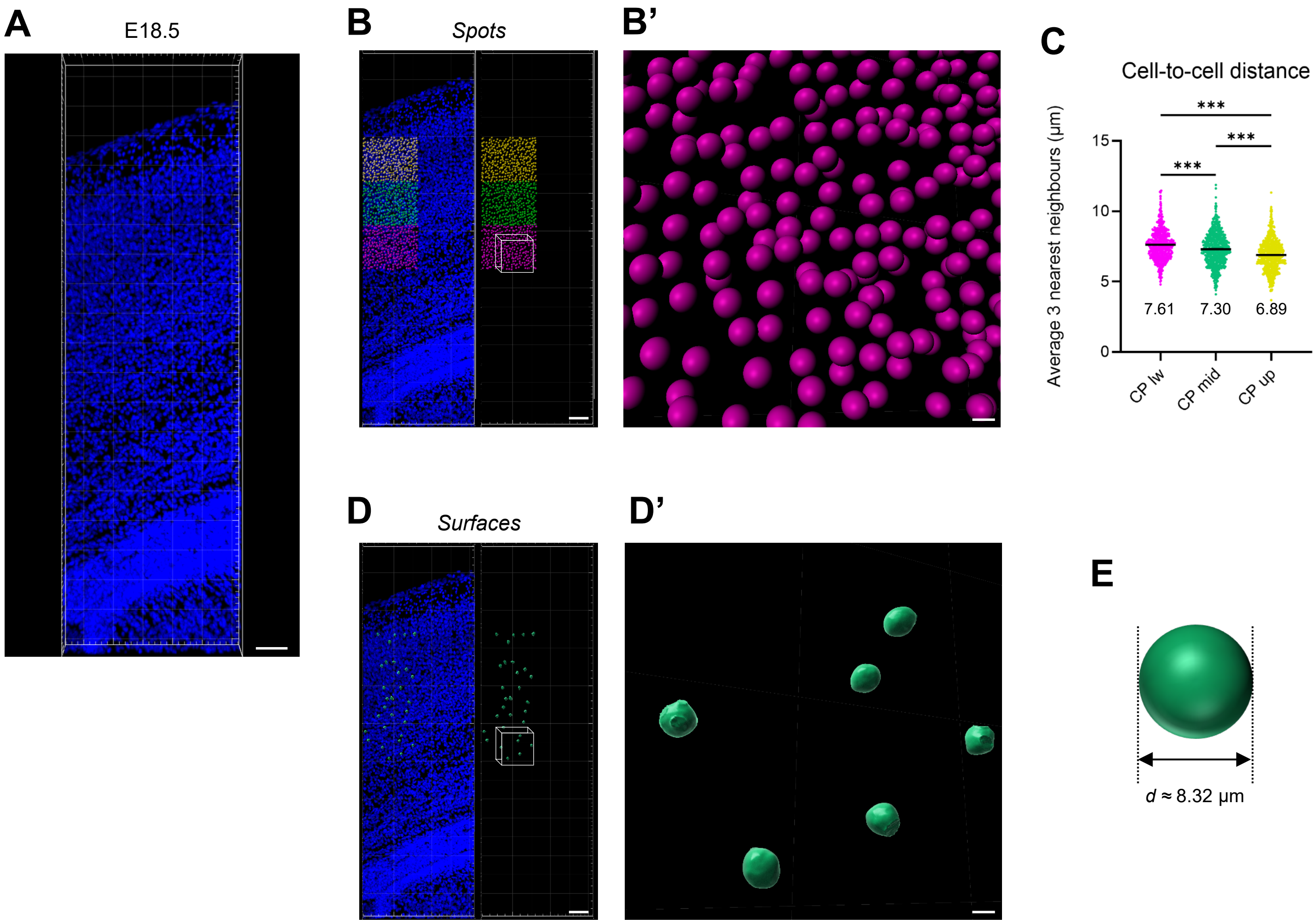


**Fig. S5. Cell-to-cell distance and nuclear volume within the CP.**
(**A**) Coronal brain sections from E18.5 mouse somatosensory cortex were counterstained with DAPI (blue) and reconstructed in 3D using Imaris.
(**B**) The dataset from (A) was analyzed using the Spots function in Imaris to measure intercellular distances in 3D. Spots were placed at the centers of nuclei based on the DAPI signal, and the X, Y, and Z coordinates were collected. The CP was divided into three equal regions: lower CP (CP lw; magenta), middle CP (CP mid; green), and upper CP (CP up; yellow).
(**B'**) Enlarged view of the boxed region in (B).
(**C**) Quantification of the mean distance to the three nearest neighboring nuclei based on the analysis in (B). Counted cells for CP lw, CP mid, and CP up were 645, 680, and 742, respectively. Mean values for each area are indicated on the graph. Statistical analysis was performed using one-way ANOVA followed by Tukey's multiple comparison test. Error bars represent the mean ± SEM. ****P* < 0.001.
(**D**) The dataset from (A) was further analyzed using the Surfaces function in Imaris to reconstruct nuclei and measure their 3D volumes. (**D'**) Enlarged view of the boxed region in (D).
(**E**) Based on the averaged nuclear volume (304.9 $\mu m^3$), the mean diameter was calculated to be 8.32 µm (n = 60 cells).
Scale bars: 50 µm (A, B, and D); 7 µm (B', D').

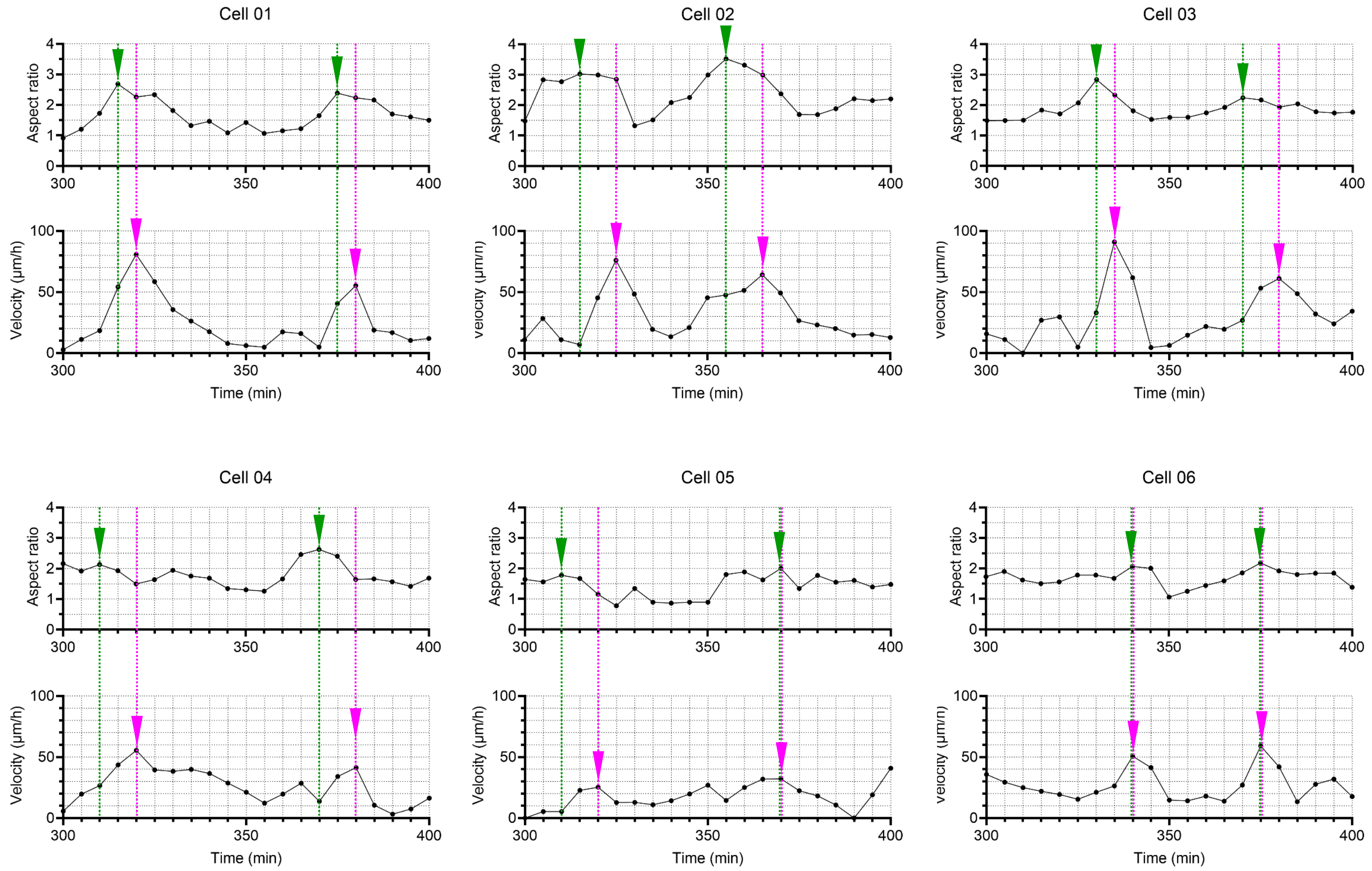


**Fig. S6. Quantification of nuclear dynamics in individual EGFP⁺ neurons in the CTRL group from Figure 5E.**
The nuclear AR (top) and migration velocity (bottom) of each representative cell were quantified over time. Green arrowheads indicate local maxima in nuclear elongation, while magenta arrowheads represent peaks in migration velocity. Dashed vertical lines connect the AR (green) and velocity (magenta) traces to highlight their temporal correlations.

**A**

| | Cell name | Family | Gender | Date of biopsies | LMNB1 | Clinical symptoms | Clinical signs | IPSCs (lines) |
|---|---|---|---|---|---|---|---|---|
| CTRL | CTRL 4 | MA | | 2011 | negative for duplication | | | 6 |
| | CTRL 5 | MA | | 2011 | negative for duplication | | | 3 |
| ADLD | ADLD CMV | CA | F | 8/9/2017 | | syntomatic | | 6 |
| | ADLD MUSI | MU | F | 2017 | | syntomatic | | 8 |
| | ADLD 1 | MA | M | | Duplication of 324,675 bp identical in all the family - (hg19) chr5: (12 6,040,794-126,365,469)x3. Tandem duplication with an insertion at t he breakpoint of 12 nucleotides (ATGTTTGTATTT) | | Brain magnetic resonance metabolic and microstructural changes in ad ult-onset autosomal dominant leukodystrophy DOI: 10.1016/j.brainresbull.2015.07.002 | 2 |
| | ADLD 2 | MA | F | | Duplication of 324,675 bp identical in all the family - (hg19) chr5: (12 6,040,794-126,365,469)x3. Tandem duplication with an insertion at t he breakpoint of 12 nucleotides (ATGTTTGTATTT) | | | 9 |
| | ADLD 9 | ZA. | F | | | asyntomatic at ti me of biopsy | | 8 |

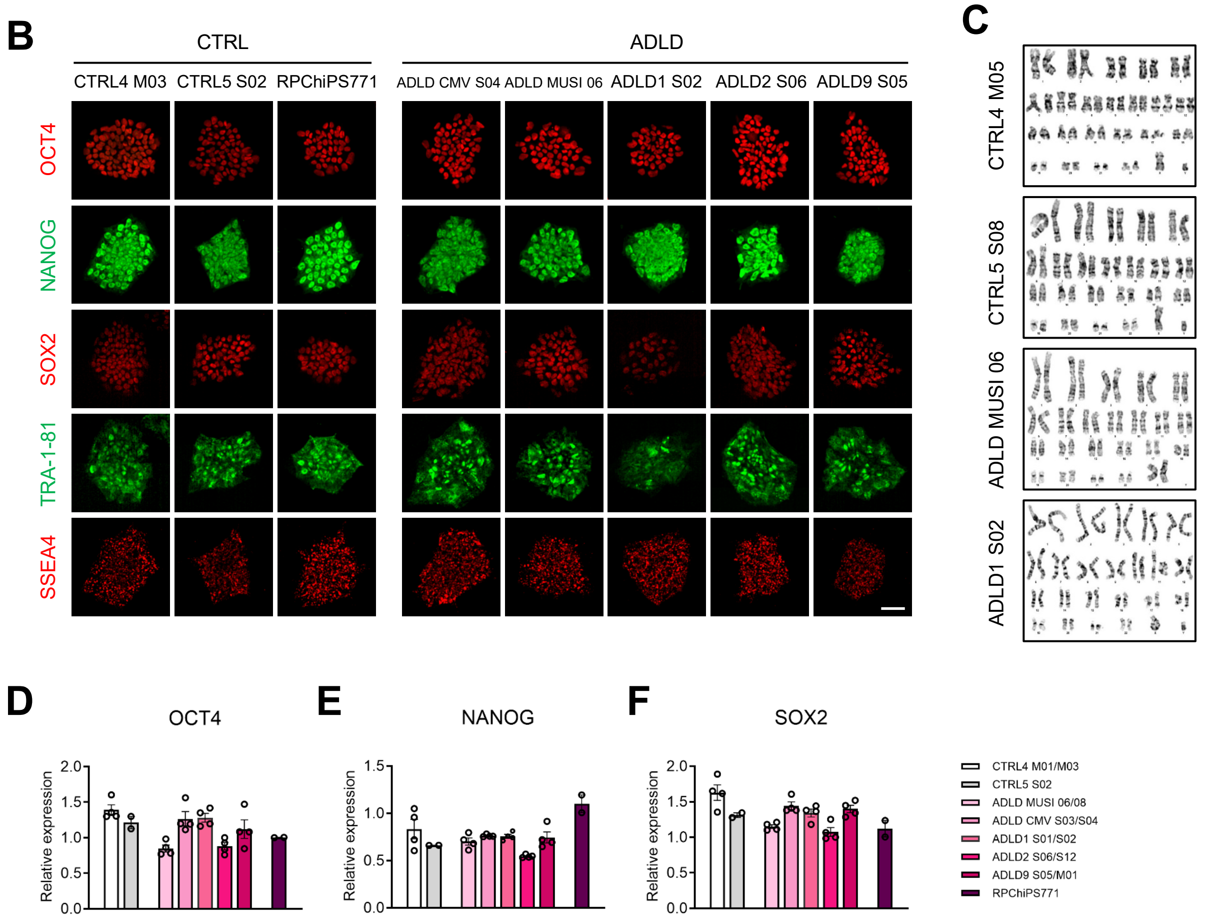


**Fig. S7. Characterizations of iPSC lines derived from CTRL and ADLD patients.**

(**A**) Clinical and genetic information for CTRL and ADLD patient samples used for iPSC generation.

(**B**) The generated iPSCs were stained with antibodies against OCT4 (red), NANOG (green), SOX2 (red), TRA-1-81 (green), and SSEA4 (red).

(**C**) Representative karyotypes of CTRL and ADLD patient iPSCs, all demonstrating normal chromosomal profiles.

(**D** to **F**) Quantification of relative mRNA expression levels of *OCT4*, *NANOG*, and *SOX2* in iPSCs, measured by qPCR. RPChiPS771 was used as the reference control (Park et al., 2022). Each dot represents an independent experiment.

Scale bars: 50 µm (B).
